# Beyond fitness: selection and information flow through the constructive steps in lifecycles

**DOI:** 10.1101/2021.02.09.430402

**Authors:** Eric Smith

**Affiliations:** Department of Biology, Georgia Institute of Technology, 310 Ferst Drive NW, Atlanta, GA 30332, USA; Earth-Life Science Institute, Tokyo Institute of Technology, 2-12-1-IE-1 Ookayama, Meguro-ku, Tokyo 152-8550, Japan; Santa Fe Institute, 1399 Hyde Park Road, Santa Fe, NM 87501, USA; Ronin Institute, 127 Haddon Place, Montclair, NJ 07043, USA

**Keywords:** Fitness, Price equation, Lifecycle, Hypergraph, Large deviation function

## Abstract

The replicator is the fundamental abstraction of evolutionary genetics. Only for replicators do Darwin’s concept of fitness as differential reproductive success, and the formalization by Fisher and Price in terms of apportionment of descendant populations to ancestors, coincide without ambiguity or potential conflict. The organization of the Price equation, causal interpretations of Fisher’s Fundamental Theorem and its relatives, and the abstraction of fitness as the sole channel through which information flows in from environments to form the adapted states of evolving populations, all follow from properties of replicators imposed artificially on the genetics of more complex lifecycles. Here it is shown how to generalize this role of the replicator to the autocatalytic flows in the generators of Stoichiometric Populations Processes, and to generalize from the unique summary statistic of fitness to a class of summary statistics that appear as regression coefficients against the autocatalytic flows associated with reproduction, including replication but also including constructive operations beyond simple copying. Both the statistical construction and the causal interpretation of Fisher’s Theorem and the Price Equation generalize from replicators and fitness to the wider class of regressions. *Ad hoc* corrections for mis-specified fitness models, which the conventional Price equation groups with “environment” effects, become part of a Fisher covariance on the basis of flows, which takes on a consistent causal interpretation in terms of an expanded concept of selection recognizing distributed information. A measure is derived for the information in the trajectory of a population evolving under a stoichiometric stochastic process, as the large-deviation function for that trajectory from a null model. The interpretation of fitness and other regression coefficients as channels for causation and information flow is derived from their inner product with the gradient of the trajectory entropy.

## I. INTRODUCTION

### A. The problem of representing biological objects

Any attempt to frame a general theory of selection in evolution [1] confronts a problem of the diversity and heterogeneity of biological objects, and of the interrelations among them. A part of the difficulty of representing the objects responsible for evolutionary change is captured in debates over *units of selection* [2–6] or *levels of selection*[7].^1^

The key derivation that enabled formalizing the concept of levels of selection was the Price equation [9–11], which clarified the status and scope of expressions for evolutionary change in terms of covariances with fitness, including the growth-rate equation and Fisher’s Fundamental Theorem of Natural Selection (FTNS) [12, 13].^2^

Important though it has been, the Price equation and all formalizations of levels or units of selection that follow from it obscure a problem in the representation of biological objects as fundamental as the problems it illuminates: the Price equation is predicated on a definition of *fitness* that only applies to a restricted subset of the events that occur within biological lifecycles.

The statistical definition of fitness by Fisher [12] and Price [1, 9] – an apportionment of objects in an offspring generation to objects in the parent generation according to the parents’ types – only coincides fully with Darwin’s concept [15] of differential reproductive success for an even narrower set of objects, which are the biological *replicators*. Only for these does the mechanics of descent assign each offspring unambiguously to a single parent at the same phase of the lifecycle. Price himself recognized how much of reproduction the replicator abstraction excludes, and generalized the statistical notion of fitness (and of selection as Price defined it in terms of fitness [1]) to a more flexible notion of “corresponding sets” that could be assigned within parts of lifecycles, where the objects (re)producing and the objects produced could be of different types (e.g. adult diploid organisms and their haploid gametes). While greatly increasing the versatility of the Price equation and its capacity to describe heterogeneous units of selection, the relation of corresponding sets that Price kept is ultimately the feature that excludes a large class of lifecycle events that should be included as channels for selection.

This paper argues that certain inadequacies of the standard formulation of population genetics to furnish a comprehensive theory of selection derive from limitations in the concept of fitness itself, and of the correspondence relation that is essential to Fisher’s and Price’s definition [1, 9, 12] of fitness. They are built into the structure of the Price equation as it is currently derived, but even more fundamentally they are built into the dependence of population genetics on a concept of the replicated gene that emerged from the Modern Synthesis [16].

These limitations are not defects in the definition of fitness, however, and the paper argues for fitness to remain as understood by Price. The problem, instead, is an inadequate representation of objects in terms of “hereditary particles”[17] (derived from the replicator abstraction for Mendelian genes), and of the role of relations within and between objects as carriers of evolutionary information.

An adequate formal theory of selection must include a model of *construction*^3^ within lifecycles alongside the more limited abstraction of replication, and an additional suite of summary statistics characterizing selective differences in construction, to augment the differences in replication characterized by fitness. In terms of these, the statistical program of Fisher and Price can be carried out as in the standard formulation of population genetics. The resulting forms not only encompass aspects of selection excluded by Price’s [1] generalization to corresponding sets; they also handle many problems of multi-level selection in a more transparent and natural way.

### B. From replication-based to construction-based abstractions of lifecycles

#### 1. Replicators are too limited to separate model structure definition from model solutions

Population genetics is built over the abstraction of the replicating gene [17, 20]. Lineages of genes abstracted as hereditary particles, however they may be combined in packages in different phases of a lifecycle, are trajectories branching *forward in time*, with each daughter particle having a unique mother particle. For genes by themselves, this forward branching defines fitness as a summary statistic directly from the mechanics of descent. It is the feature retained in Price’s [1] corresponding sets.

The limitations of the underlying model are reflected in the way the Price equation is written for lifecycles in which packages as well as genes have fitnesses affecting overall reproductive success. (Dominance and epistasis are examples of fitness inherent to “packages”.) The FTNS - the expression for what Ewens [21, 22] terms the partial change in characters arising from their covariance with additive fitness - assimilates diploid reproduction, a process that does not branch only forward in time, to Price’s corresponding sets by attributing only the additive genetic value or the breeding value [14] of offspring characters including genotype or fitness to each of its parents. (The notion of packages is not restricted to mean organisms, though the original application of the FTNS was mainly concerned with this case; the indexing and algebra of the Price equation apply equally naturally to events jointly generated by sets of organisms [23], regarded as a kind of “phenotype” of the collective.)

What the Price equation then represents as “descent”, which should define a structural property *of* the lifecycle model, depends in the FTNS on the values of regression coefficients that are solution properties *within* the model. The many problems of interpretation of the covariance with additive fitness [14, 22, 24], and the ambiguities in what constitute units of selection, ultimately derive from the lack of a well-defined separation in this concept of descent, between the structural definition of models and the properties of solutions within them.

#### 2. Stoichiometric Population Processes as an alternative abstraction

In this treatment we therefore dispense with the abstraction of the replicating gene at the outset, and adopt a different formal model for biological lifecycles. Genes are not gone from model parametrization; they may be present as quantities conserved within parts of lifecycles (Sec.V A), or they may be the basis for rules to construct objects and interactions (Sec.X C). What is important is that counts of genes do not define the whole potential of objects to carry an evolutionary state through time.^4^That additional freedom in the underlying representation is needed to avoid the limitations inherent in representations defined by composition of replicators.

The abstractions adopted here for lifecycles will be called *Stoichiometric Population Processes* (SPPs). They are used in many domains under different names, to represent chemical reaction networks [25–28], or as dynamical models of process execution in computer science (with some further refinements) under the name *Petri nets*[29]. SPPs employ a heterogeneous object model. The state of an evolving population (in the Markov sense) is carried by the counts of individuals classified into *species*, which may be of diverse kinds. In this respect SPPs admit the same generalization as Price’s corresponding sets.

SPPs then take one step further, which is the natural completion of Price’s [1] insights. They are not limited to population changes in which one object is replaced by one or more descendant objects in a corresponding set. Instead, the elementary events of population change, which we term *reactions* following the usage in chemistry, remove sets of individuals, and deposit other sets into the population in their place. The shift from corresponding sets to general set-wise transformations replaces an ordinary graph representation for lifecycles with a construction called a *hypergraph*[30], described further in Sec.II C. The set compositions of the inputs and outputs of each reaction are termed the *stoichiometry* of the reaction, after which SPPs are named. The species types, and the hypergraph of allowed reactions with their stoichiometries, define the lifecycle model in an SPP.

SPPs can represent a variety of modes of construction within complex lifecycles, including but also extending beyond acts of replication. This paper will be concerned particularly with *autocatalytic flows*[31], which are derived from the topology of the hypergraph representing a lifecycle. These are sets of coupled transformations that connect equivalent phases within lifecycles in the same way that replication does, without requiring continuity of any single type of object throughout the flow. A single flow may be mediated by diverse types of objects and by sets of objects. The statistics from regressions against a basis of flows therefore express relations that make biological objects more than, or different from, mere lists of their parts. The higher-order statistics of flows characterize selection with a completeness that fitness alone generally cannot.

It is important in what follows to distinguish regression on autocatalytic flows from two constructions within standard population genetics that look similar notationally and incorporate parts of the same information. The abstraction and generalization of a gene or organism identity on which population dynamics might depend is what Frank [17] terms a *predictor*. It is clear that an event of population change might be generated from an interaction of entities – whether haplotypes in a diploid organism or a host and a pathogen during an infection – which are joined in the interaction but not in heredity. Therefore, set-valued predictors should generally be used in the regressions that partition fitness and environment terms in the Price equation. The outcome generated in aggregate may even be modeled as a phenotype of the set [11], allowing parts of evolutionary change that a Price equation for frequency-dependent fitness of individuals would place in the environment term to be moved into the covariance of the Price equation. However, as long as some Fisher/Price apportionment must be used to represent the lifecycle within the Price equation, at most breeding values for fitness or other characters may be expressed in the covariance term, and a well-defined separation between process description and model solution remains broken. These limitations, imposed artificially by apportionment on lifecycles that lack them in reality, are not imposed by flows on the hypergraph.

### C. Causal interpretation of covariances

Evolutionary genetics is meant to be a causal account [17] of the long-term change in populations produced by selection. In particular, the covariance terms in the Price equation (the growth-rate theorem, FTNS, Robertson’s secondary theorem in quantitative genetics [32, 33], etc. [11]) are widely regarded and presented^5^ as separating out the parts of a change in population state “due to” selection [10], or “due to changes in gene frequency”[14]. Covariance with fitness is meant to contrast with other sources of change such as altered environment or character changes during transmission such as occur through mutation.

Failures of covariance terms to provide a causal account of evolutionary change appear as failures of *dynamical sufficiency* [10, 34]. A descriptive account of the change over one generation or one interval of continuous time in the Price equation fails to furnish model coefficients that predict the population behavior at other times, and a regress of higher-order moments must generally be solved to obtain a predictive model [35, 36].

The analysis below will draw a sharp contrast, between the Price equation and the interpretation of covariance with additive genic fitness, and the corresponding constructions over autocatalytic flows in an SPP. In the former construction defined by reduction to replicators, the term in the Price equation described as representing “environment” or “property” changes, in fact generally subsumes a variety of *ad hoc* corrections for the mis-specification of additive fitness as a dynamical model. In the Price equation over flows in an SPP, the same contributions to population change are shifted into the covariance expression, where they take on a proper interpretation as changes due to selection.

The difference of this construction from its nearest counterpart in the conventional approach, with a set-valued phenotype and set-valued predictors, is that in the latter only the breeding values are shifted from the environment to the covariance term [11], generally leaving a non-vanishing environmental contribution, and a covariance term that on its own is not dynamically sufficient. In the regression on flows, terms that Price would have categorized as redistributions appear naturally in the covariance with no residual environment corrections.

### D. Information and information rate for an evolving trajectory

Dynamical sufficiency provides a criterion for whether a model estimated from regressions captures all the information in a population imparted by selection. The stochastic formulation of SPPs defines a *large-deviation function*[37, 38] for any observed population trajectory [39, 40] from the trajectories typical under a generative model. That function is put forth here as a measure of the information “in” the population trajectory left unexplained by the model. Minimizing (over model parameters) the information left unexplained in an observed trajectory generalizes common least-squared-error criteria from statistics.

Error criteria defined over entire trajectories provide the dimensionality needed to estimate the larger array of parameters offered in SPP models which would be underdetermined at a single time, a consideration of statistical power that could otherwise favor projecting down onto low-dimensional but mis-specified additive models. In certain limits of slow population change, where the whole-trajectory information converges to a time-integral of a local function of the rate of change, Fisher’s information metric is recovered in the rate of input of information by selection. The basis of autocatalytic flows on which the regression models are computed allows a definition, from the large-deviation function, of channels for information flow into the population state from different factors of selection.

### E. Organization of the presentation

The presentation of the material is organized as follows: Sec. II fills in some detail omitted in this brief introduction, about the way genetics casts evolution solely in terms of changes in frequency of genes or genotypes, the role of replicators underlying this characterization, the problems of consistency in such a formulation even within the conventional arguments, and how some information about what genes *do* must be part of genetics to overcome these defects and inconsistencies. The section is concerned with the logic that makes a definition of selection in terms of fitness inherently insufficient, and with the SPP as the natural remedy. Readers who wish to see immediately how stoichiometric models are built and how they are used in examples may wish to skip Sec.II in a first reading.

Sec. III introduces the simple class of diploid lifecycle problems that will serve as a running example of methods throughout, first as discrete-generation Wright-Fisher models and then in continuous-time forms that are simpler under the solution methods used here. The models used will be population-genetic, with “identified genotypes”[41] and fitness or other characters taken to be indexed by these, for ease of presentation. The limitation to additive characters and breeding values remains the same in quantitative genetics with genotypes not explicitly identified [23], as the origin and structure of the Price equation remain the same [11], but a formulation in terms of quantitative phenotypes is not pursued here.

Sec. IV introduces hypergraph representations for the lifecycles that generate SPPs, emphasizing the different roles of graph stoichiometry versus rate constants.

Sec. V introduces autocatalytic flows on a hypergraph, and shows how regression coefficients in the basis of flows generalize Fisher’s definition of fitness from regression coefficients in the basis of replicators.

Sec. VI decomposes the rate equation for population change under the reasoning used by Fisher and Price, first in traditional terms of additive genic fitness plus an “environmental” remainder, and then in the basis of autocatalytic flows. For the examples here, a distinct environment term vanishes altogether from the latter, as all effects of both viability and the mating system are organized into covariance expressions in the form of a growth-rate theorem. **The resulting equation (42) is the first main result of the paper**.

Sec.VII uses the Price equation in the autocatalytic flow basis to study two regimes of selection involving biased reproductive steps. In the first, with context-independent asymmetric gametogenesis, the Price equation for flows decomposes into a sum of exact genic and genotype fitnesses. In the second, production of haploid gametes and survival of diploid adults are made conditionally dependent, giving an example of the contrast between within-level and between-level selection.

Sec.VIII introduces the large-deviation function for the trajectory of an evolving population as a measure of information imparted through evolution. **A general form (77) and its approximation (80) for slow rates of change are the second main result of the paper**. The Fisher metric arises naturally in the rate of information incorporation, along with a useful construction for the sampling rate that replaces the role of whole-population size in defining covariance in discrete-generation models.

Sec.IX again studies selection regimes, this time in relation to trajectory information measures. Estimation of generating models on trajectories provides the statistical power to identify the additional parameters in an autocatalytic-flow model that would be confounded in regressions at a single time. Channels for information flow are then exhibited, and “causes” of population change in the growth-rate theorem are sorted into sources either of information or of randomness.

This paper uses the simplest non-trivial examples throughout to introduce core concepts with minimal distraction to model details. Much of the power of stoichiometric modeling, however, comes from its versatility and its capacity to organize problems in which complex and structured collections of types may be needed to understand evolutionary change. Sec.X discusses richer applications and additional concepts pertaining to model complexity.

## II. WHAT IS EVOLUTION?

Genetics is on one hand just an accounting system defined over the abstraction of replicators (and for some applications fitness); on the other hand it is a convention for projecting the vast complexity of life down onto that accounting problem, which claims to furnish a representation of what evolution “is”. Before offering an approach to genetic problems that starts from any other abstraction than the replicator, it seems necessary to clarify what conceptual divisions the current approach entails, and to argue that some of biological function currently placed outside the scope of selection, genetics, and the essence of evolutionary change, in fact belongs within the formal treatment of those concepts.

### A. Changes in frequencies of objects

Prior to and superseding any disagreements about levels or units of selection, a shared commitment of all approaches to population genetics is an abstraction of evolution as the change in frequencies of some collection of objects [42]. The reduction of evolutionary change to an account of frequencies depends on two premises:

1. that it is sensible to consider the frequency of objects separately from questions about what those objects *do*. The latter are left as problems for other domains of biology, and their solutions feed back into genetics only through the map of phenotype to fitness or other characters;
2. that objects, the durable entities carrying the population state in the Markov sense, are distinct in that role from relations among objects, and that the objects are sufficient. The relations do not carry memory (and thus are not targets of selection) because they are dissolved and re-created in different forms rapidly through time [43, 44].

The question: in terms of which objects can evolution be reduced to a counting problem, clearly depends on judgments about what kinds of approximations are acceptable and which distinctions matter. For instance, when should an organism with a whole genotype, or the entire sequence of a chromosome, be treated as an object carrying the memory of the population state, and when is either of those at most a collection of lower-level objects passing through an ephemeral relation?

### B. Mendelian heredity and the importance of replicators to the conception of fitness

The reduction of all long-term evolution onto the counts of genetic objects must be understood in relation to the other dominant abstraction in genetics: of replication and replicators. Replication, in contrast to the more general concept of reproduction, entails a direct act of copying. Projection of evolution onto replicator models limits the representation of information to that for which a template/copy relation is explicit.

The Modern Synthesis was a project to instantiate Darwin’s pre-formal requirements for evolutionary heredity on a foundation of Mendelian genes [16]. The goal of quantifying fitness in a manner compatible with Mendelian heredity led to Fisher’s definition in terms of apportionment of offspring to parents, which serves the very important function of separating the problem of statistical reduction of diachronic histories from that of statistical inference of causal models [17] of the generators for such histories,

Both the limits on the summary statistics used in reduction, and the limits of complexity in causal models, are choices about what information a state can carry or transmit. The dynamical sufficiency [34] of a model gives a measure of how much of selection and heredity is captured with those choices. Strict genic selection treats a composite entity like a chromosome as nothing more than the list of its genes, while a model of the chromosome as a unit of selection [2, 34, 45] casts the full sequence as a primitive object that can be transmitted.

The segregating allele and the discrete copying operation are highly abstracted and coarse-grained patterns projected onto complex and multistep constructive processes through which living matter is assembled and operates. Nonetheless, the 1953 discovery of a molecular mechanism for DNA copying [46, 47] reified the Mendelian statistical gene of [48] in matter as a replicator. Examples of this point of view are such formulations as [49]: “It is useful to begin by defining the word ‘gene’ to mean a particular copy of a nonrecombining sequence at some locus in some individual.”

The tension that results from insisting that the replicator abstraction defines what evolution “is” in terms of counting, and the inability to commit categorically to what qualifies as a replicator, is evident even in sophisticated treatments. Lynch [42], for instance, in arguing that relational information is not heritable in the long term and therefore cannot be needed to define evolutionary change, nonetheless closes with

> “Organisms are more than the sum of their parts, just as genes are more than the sum of their functional components. But if we are concerned with the process of evolutionary change, then evolution is indeed a change in *genotype* frequencies.” [emphasis added]

Sets of genotypes differ from sets of genes only in co-occurrence statistics, which are the relational variables whose status as targets of selection is under dispute.

The need to satisfy incompatible claims for what can be transmitted motivates Price’s separation [1] of lifecycles into stages, with objects at different levels of organization transmitted within different stages. Although no longer replicators by the criterion of providing continuous templates across entire lifecycles, Price’s corresponding sets were chosen to retain his own [9] and Fisher’s definition of fitness in terms of apportionment.

### C. Construction: making objects more than, and different from, collections of parts

Mendelian heredity as a constraint on lifecycles in no way implies that information is modularized within replicators, or that template/copy relations exist for most biological characters. Information can be, and often is, carried in distributed form across collections of objects.

Patterns transmitted through time by distributed instructions include the complex objects and transitions that constitute stages of lifecycles. These, like replication events, emerge in the aggregate from a complex, constructive choreography of microscopic events; they differ only in the absence of a physically persistent template object connecting a stage of the lifecycle in an ancestor’s generation to the same stage in the descendant’s generation.

Redistributive steps, such as fertilization or recombination, introduce mixing timescales for the memory of some variables, while others characterizing the lifecycle are unaffected. Chromosome structure and ploidy, the degree of self-compatibility or incompatibility in mating, and other features affecting long-term evolution breed true to type.

The persistence of lifecycles as traits, moreover, is as “genetic” in its origin as the variation of traits modularized within alleles: lifecycle traits can vary among species, and are protected from corruption or mixing by mechanisms ranging from restriction/modification systems to inter-species infertility or hybrid sterility. The architecture of the lifecycle is transmitted accurately within the group, at the same time as other traits that factor into contributions from genes can arise in combinations that do not outlast individual organisms.

The robust use of distributed information requires that, in addition to the frequencies of template and copy “objects”, the kinds and frequencies of the organizations in which they occur must be memory variables and targets of selection. The aspects of the phenotype that control these organizations are the aspects of “what genes do” that belong in a more expressive abstraction for population genetics.

As mentioned in the introduction, Frank [17] recognizes that, while any Price equation may give a “complete and exact”[50] descriptive account at a single time, the basis of “hereditary particles” in which additive fitness is defined will not generally provide the most complete set of predictors possible for causation. Set-valued predictors may improve the dynamical sufficiency of the resulting statistical description. Sets provide a representation of one kind of relation among elements (though they stop short of the further abstraction of relations as rules discussed in Sec.X C). Price’s [1] construction of corresponding sets recognizes the need for a different kind of generalization, beyond representing biological objects as sets or lists of components, so that objects of arbitrary kinds may stand on their own as distinct types. The full potential of this heterogeneous object model is emphasized by Queller [23] in attributing phenotypes to sets of organisms defined from their interactions.

The stoichiometric representation of lifecycles used here makes objects more than and different from sets of other objects by changing the way objects predict dynamics from the abstraction of Price. Under the ordinary, forward-branching graph defined by Price’s corresponding sets, the objects that terminate a graph edge both carry the population state and directly index its transformations. The hypergraph of the SPP, in contrast, distinguishes the species carrying the population state from the input and output sets that terminate a hyper-edge (termed *complexes* [25, 26]), which govern state transitions. Species are persistent, whereas complexes are merely samples with no persistence of their own. In SPPs, a constructed species is different by default from any collection of inputs that may (*via* some reaction) determine its type.

The mathematics and interpretations of SPPs are extensively developed for the representation of chemical reaction networks (CRNs) [25–28, 51–55]. They will provide natural expressions for the generators of biological lifecycles not because organisms and their components are the same as chemicals, but because certain problems of representing objects and relations are common to both. In CRNs the objects are molecules and conversions among sets of objects are reactions. A set of components (atoms or molecular fragments) may be combined in some new molecule, but the new molecule is distinct from the list of its components in that it also incorporates constructed relations among them (bonds), and thus new molecular fragments that were not part of any input molecule. (See [56, 57] for category-theoretic methods to express these relations.)

For lifecycles, rather than imposing correspondence relations on the state in the form of sets as in [1], we may take each type of individual: gamete, zygote, sporophyte, gametophyte, etc. to be a state-carrying object of its own type, allowing the stoichiometry of the transitions to handle set correspondences. The way Mendelian genes may be recovered from conserved quantities of the stoichiometry of subgraphs is illustrated in Sec.V A, and more about their use in rules is discussed in Sec.X C.

A representation of lifecycles using hypergraphs differs in one important respect from the construction given in [58], which clearly recognized and aimed to address the same limitations of the genetic representation that we address here. Kerr and Godfrey-Smith generalize Price’s corresponding sets from a single forward-branching network to a superposition of such networks, each representing a distinct source of inheritance. From these they derive a pair of covariance terms in a generalized Price equation, governing the forward-branching and backward-branching relations that their construction allows. The conceptual difference between a hypergraph and a sum of simple graphs is that events in the hypergraph impose *concurrency* on the conversion of the input set into the output set [59, 60]. The expressive power of hypergraphs is attested by the fact that many problems of search and optimization that reside within simple computational complexity classes on ordinary graphs become computationally hard (NP) on hypergraphs [61]. (See [62] for a thorough introduction to computational complexity classes.) While the kinds of problems typically considered in population genetics will not be computationally hard, the *capacity to represent such hard instances* characterizes the expressive flexibility of the abstraction.

### D. Evolution and selection beyond replicators and fitness

SPPs offer an alternative compact abstraction onto which to project what is essential about evolution. In place of characterizing evolution as the change in the frequency of replicators (or of some *ad hoc* collections above the level of replicators, such as genotypes), we can identify evolution with *the change in population states through time by the recurrent passage through lifecycles*.

In starting from lifecycles rather than genetic lineages, we still require regularity: unstructured change without a recurring reference stage would be too general and should not qualify as evolution. Representing different states in a lifecycle as independent objects simply reflects the reality that a gene always exists in some context: in a plasmid, chromosome, *n*-ploid genotype, or potentially higher-order structure.

Lifecycles need not be regular or rigid in the way replication events are: complex lifecycles [63], with multiple free-living phases, structured populations leading to competition at many levels, and environmentally conditioned transitions between phases, fall naturally under this category (though they are not pursued in this paper).

The concept of selection that Price sought in terms of differential survival or replication in corresponding sets generalizes to one based on differential rates for autocatalytic flows through lifecycles. Price [1] observed that: “Selection requires variation and acts to reduce variation: this results in a competing relationship between selection and entropy increase about which it would be interesting to have deeper understanding.” The large-deviation measure of entropy in Sec.VIII provides systematic ways to understand that relationship.

The defining property of large-deviations scaling [37] is a separation of scale from structure [38]. Total population size, which may expand or contract, serves as the scale factor in the absolute entropy or potential information in a trajectory. The other factor, associated with structure, is called the large-deviation *rate function*. It is a specific entropy that measures the diversification in the population. The derivatives of the large-deviation function with respect to each flow in the lifecycle determine how their rates interact with the population state to increase or decrease either the scale or the diversity contributions to entropy.

## III. SIMPLE DIPLOID REPRODUCTION MODELS TO ILLUSTRATE THE MAIN IDEAS

The approach in this paper will not be to develop a general notation and system for stoichiometric modeling of lifecycles (a project for other work), but rather to develop a simple running example that is first modeled conventionally in terms of fitness, and then in the more complete stoichiometric breakdown. The emphasis is on decomposing reproductive success in terms of the independent flows in lifecycles, in place of Price’s dichotomy of corresponding sets with intervening redistributive steps. The model is chosen for minimality and considers only dominance. The effects demonstrated generalize directly to epistasis, but problems related to combinatorial complexity are not addressed.

### A. Conventional Wright-Fisher discrete-generation models

The example is a single-locus, two-allele system for a sexually reproducing diploid, drawn from [14]. Mating types of gametes, and sexes of diploids, will not be distinguished. Gametes will not be treated as free-living, and gametogenesis and fertilization will serve simply to multiply and redistribute alleles. The numbers of diploid organisms will thus determine counts of both genes and genotypes. Within that standard background lifecycle, two limiting forms of fertilization will be considered: random mating in the population and self-fertilization. Intermediate degrees of self-incompatibility can be treated with mixtures of random-mating and selfing models.

For reference to standard treatments [64], we introduce the system in a discrete-generation Wright-Fisher model. For a one-locus model, alleles and haplotypes are equivalent. Denote these by {a, A}. Diploid genotypes without order-dependence are denoted *g*∈{*aa, aA, AA*}.A state of the diploid population is indexed by a vector *n*≡(*n_g_*), where component *n_g_* counts the number of individuals with genotype *g*. Total population size is denoted 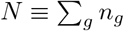. The counts of haplotypes are denoted *n_i_* with *i* ∈{*a, A*}.

Using Price’s prime notation to relate numbers in offspring and parent generations, the offspring numbers *n*’in a Wright-Fisher model with random mating and without mutation relate to parent numbers *n* as

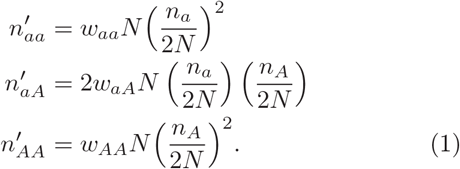

For selfing the corresponding relation is

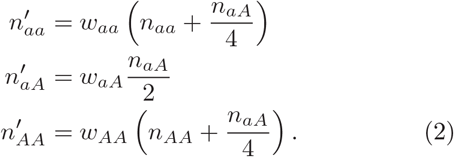

Following [14], a model of “viability selection” is assumed. *n_g_* and *n_i_* are counts at the time of mating in the ancestral generation, and 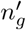 are counts at the same point in the descendant generation. *w_g_* is the fitness of genotype *g*, reflecting uniform fecundity and variable survival.

### B. Continuous-time asynchronous-reproduction processes

For the methods used below, continuous-time processes are more tractable than discrete-generation models, so we replace Eq. (1,2) with a continuous, asynchronous reproduction model that is statistically equivalent for the purposes here.

The trajectory of a population *n*(*t*) under continuous, asynchronous reproduction is represented at the level of rate equations by its time derivative, denoted *ṅ*. Beginning as in Eq. (1,2) with viability selection, the rate equations for random mating are

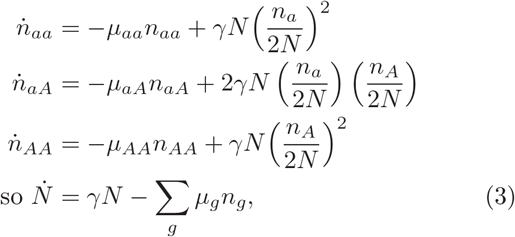

and for selfing

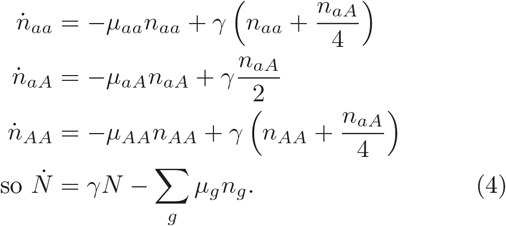

Here {*μ_g_*}are mortality rates for genotypes *g*, and *γ* is a rate of gametogenesis, equal for all genotypes *g* and alleles *i*.

The survival probability for any individual of type *g* is exponential in *μ_g_* times its age. Therefore average lifetime for type *g* is 1/*μ_g_*, and 1/2 the expected number of gametes produced during its lifetime defines an unambiguous fitness from mortality and fecundity as

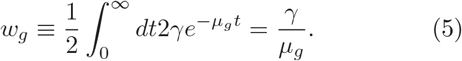

The extrapolation of the tangent of the trajectory over one such generation is the measure of population change for the continuous process that compares to the discrete-generation case. For random mating,

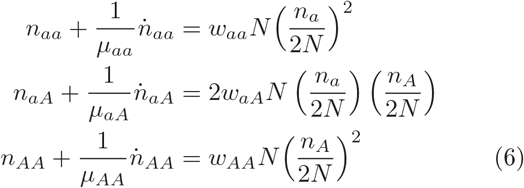

recovers Eq. (1), and for selfing,

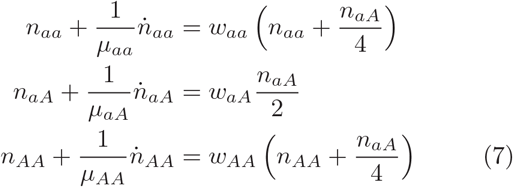

recovers Eq. (2).

The exact population change ∫ *dt ṅ* over any interval is of course different from the extrapolations (6,7), and the continuous-asynchronous model does not produce all the same statistics as the discrete-generation model, just as Wright-Fisher and Moran processes, or other canonical models, do not produce all identical statistics [64]. The purpose in these examples is to illustrate general features in a well-defined model; a considerable array of tools exist (see discussion in [40]) to incorporate into a stochastic model the many kinds of temporal or demographic structure that may characterize any particular natural population, at the cost of more elaborate analysis. A geometric picture of the invariant families associated with the two fertilization systems, and certain standard results for additive regression models, are reviewed in App. A.

## IV. USING HYPERGRAPHS TO MODEL LIFECYCLES

A consistent abstraction should separate the representation of a population process into three levels. The first specifies the parametrization of the process, which here is the *topology* determining the possible interconversions of objects. The second assigns a *rate structure* for a given case; these are the parameters for which regression coefficients provide sample estimators in a statistical analysis. Third, one may solve the process either stochastically for a distribution, or with deterministic rate equations for first moments.

### A. Topology of the hypergraph defines the stoichiometry of the lifecycle

A hypergraph is specified [25–27] by its set of *species*and its set of *reactions*. Each reaction withdraws a multi-set of species^6^from the population, and introduces some other multiset in its place. The multisets withdrawn or added by reactions are termed *complexes*.

For this example two levels of species must be defined: diploid organisms indexed by genotype *g* and gametes indexed by haplotype *i*. A third type, the null object, is introduced to allow removal of members from the population.^7^ The three classes of reactions are removal of diploids (death); gametogenesis, taken here to preserve the parent; and fertilization fusing two gametes.

Fig.1 gives a representation of the sexual lifecycle, with random mating and selfing represented in separate connected graphs. Species *g*∈*{aa, aA, AA}* and i ∈{*a, A*}are represented with solid circles. Numbers *n_g_* are state indices counting diploids. Because topology alone does not dictate whether gametes persist in numbers greater than zero, we introduce counting variables 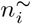 for gametes as well. (The designation *n*^~^ is adopted because kinetics will be chosen that make them ephemeral.) The null species, denoted Ø, is not associated with a counting index.

**FIG. 1:**
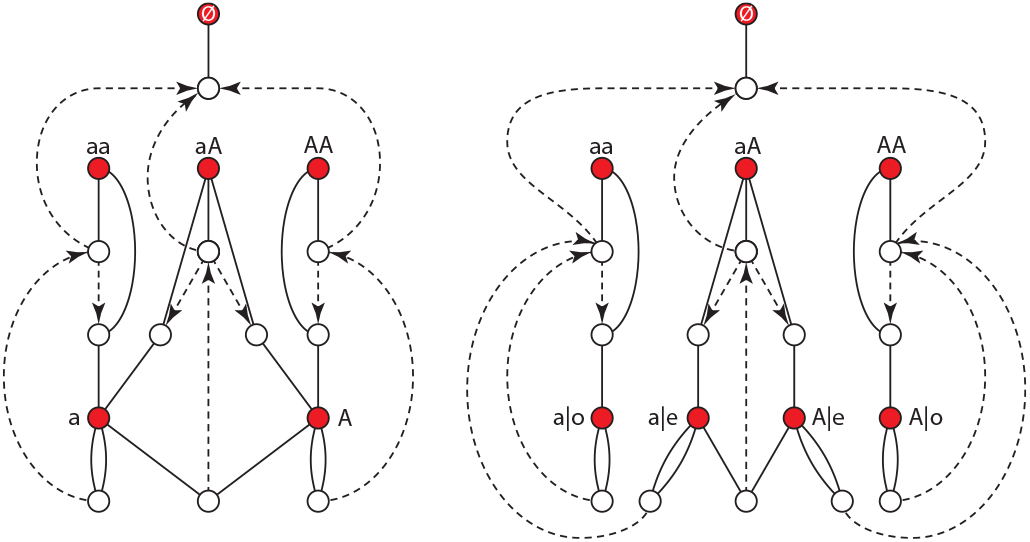
Hypergraph representations of random mating (left) and selfing (right) lifecycles. The Markov state is carried on the solid circles including diploids with genotypes *g*, gametes with haplotypes *i*, and the null object Ø. Reactions for mortality, gametogenesis, and fertilization are dashed lines with arrows converting input complexes to output complexes (open circles). Solid lines indicating stoichiometry count the number of species of each type making up a complex.

Complexes that mediate changes in the population state are represented with open circles. Each reaction corresponds to a directed dashed line between two complexes, converting its input to its output. The collection of individuals from each species that make up a complex is indicated with solid lines connecting the open and closed circles, which denote the stoichiometry of the population process. Solid lines are not directed, as a complex may be the input to one reaction but the output of another. A reaction, coupled with the stoichiometries of its input and output complexes, corresponds to an ordered pair of sets, called a *directed hyperedge*. The resulting network is called a *directed hypergraph*[30].^8^

### B. Generator graphs with rates

A hypergraph defines a generator for a stochastic process once we assign rate constants to the reactions and a sampling procedure to the complexes. Here we adopt proportional sampling without replacement, which in CRNs results in *mass-action kinetics*. Fig.2 shows an assignment of rate constants to the two reproduction models of Fig.1. Gametogenesis and fertilization are independent reactions in the graphs, and require independent rate constants. (The graph labels allow asymmetric rates of gametogenesis for the two haplotypes, but we leave these rates equal to keep algebra simple through the next few sections of this introduction.)

**FIG. 2:**
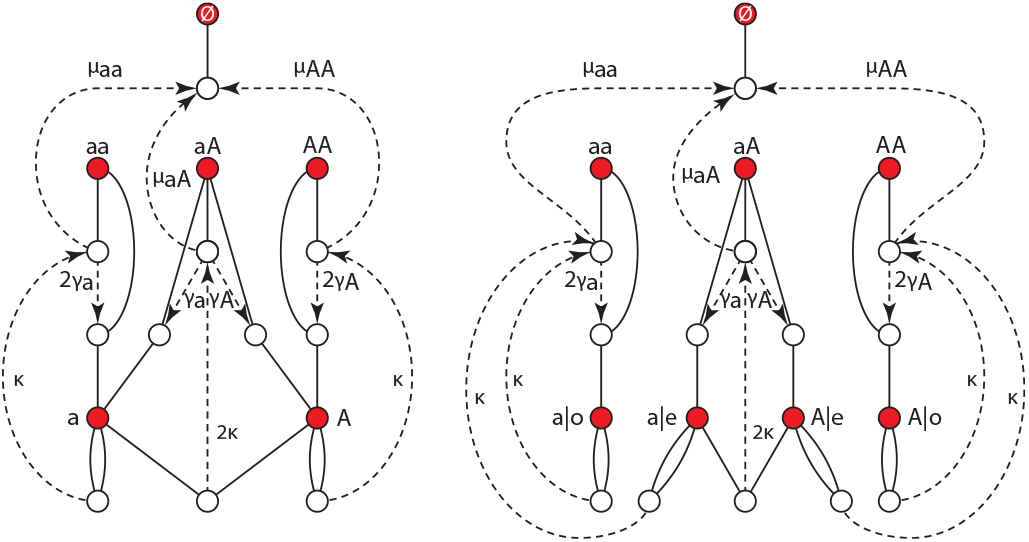
Lifecycle graphs from Fig. 1 with rate constants. *γ*_i_ are rates of gametogenesis, potentially unequal between alleles. *μ_g_* are mortality rates for diploids. *κ* is the fertilization rate.

At the level of rate equations, one can solve for the rates of change of population types with each mating system:

#### 1. Random mating rate equations

For random mating, the rates of change of gamete counts result from the net of gametogenesis and fertilization. For the two counts separately and their sum,

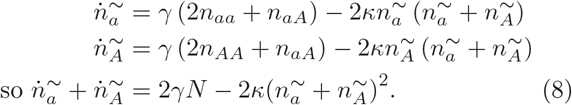

Now we introduce a limit for rate constants in the model that removes the extra independent equations brought in by the lifecycle graph relative to equations (3). Non-free-living gametes exist for periods much shorter than the lifetimes of the diploid organisms that they produce and that produce them. We can recover the statistics of this general regime of processes by taking the fertilization rate *κN* ≫ *γ*. In that limit, 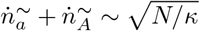 (which is 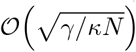 relative to the other terms in the rate equations (8) and can thus be driven toward zero) and the populations then satisfy to leading order

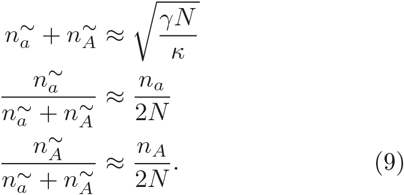

The rate equations for diploids are the net of fertilization and mortality, given by

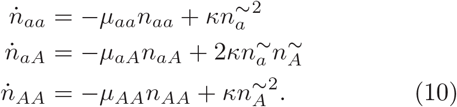

Using the approximation (9) in Eq. (10) recovers Eq. (3) to zeroth order in *1/κ*.

#### 2. Selfing rate equations

Under selfing, a more complex set of gamete types must be denoted, carrying both the haplotype of the gamete and the homo- or heterozygosity of the diploid that produced it. We denote these types respectively *i* | *o* or *i* | *e*. The rate equations for gamete populations become

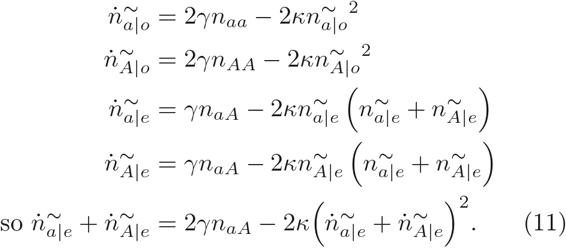

Again, at large *κ*,^9^both population size and rate of change are smaller than the input and output rates of which they are the net, resulting in

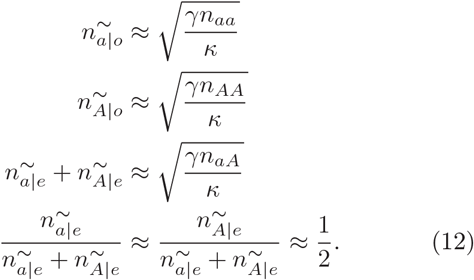

The rate equation for diploids, from fertilization and mortality, is

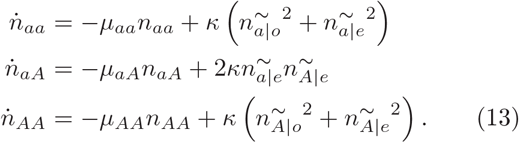

Again the approximation (12) within Eq. (13) recovers Eq. (4) to zeroth order in 1/*κ*.

## V. AUTOCATALYTIC FLOWS AND REGRESSIONS AGAINST THEM

The two processes governing change in population state are death and reproduction. In this example, mortality is a simple linear function of the population state, not requiring anything beyond the usual treatment of viability selection [14] by Fisher’s theorem and the Price equation, so we leave it aside until Sec.VI. Reproduction is the process with nontrivial structure requiring a novel treatment that the hypergraph representation of the lifecycle provides. This section develops the mathematical analysis as a generalization of the analysis of replication.

Fig. 3 shows a degenerate lifecycle model that treats diploids as replicators. Such a graph might be adopted as a null model if one were faced with a population of organisms about which nothing was known of the genetics or reproduction system, and all these were to be inferred statistically from its population trajectory. (These null models provide one way in Sec.VIII to define the “total” information that a population trajectory reflects, not only about differential mortality and fecundity, but about the lifecycle itself as a genetically-determined cause of population change.)

**FIG. 3:**
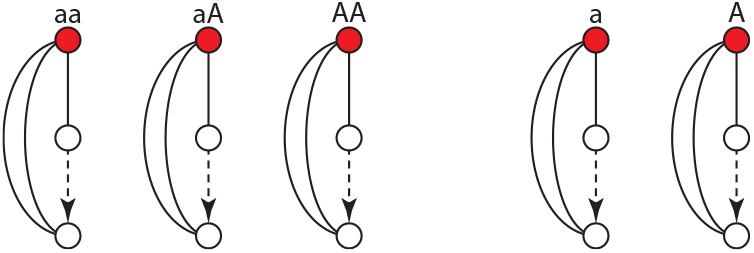
Autocatalytic flows for a replicator model acting on diploids. Each event samples some individual of a type *g*, and puts two individuals of the same type *g* into the population in place of the sampled individual.

The only reactions depicted in Fig. 3 are replications, each with the effect *g*→2*g* for some genotype *g*. Such events are the simplest forms of *autocatalytic flows* on the hypergraph: they are linear combinations of the reactions that increase the number of some species without decreasing the number of any species. The linear space of autocatalytic flows on a generator hypergraph is the domain to which we will generalize the statistical analysis of Fisher’s theorem and the Price equation.

### A. Sub-graphs with conserved quantities as a basis for autocatalytic flows

In the graphs of Fig. 1, gametogenesis is the only reaction capable of increasing the count of some species without depleting any species. To compute the effects of this single amplifying reaction on the entire lifecycle, we must first identify the basis of flows completing the lifecycle that preserve species counts within the remainder of the graph. These are identified by extracting sub-graphs of the full generator that possess conserved quantities – left null eigenvectors of the stoichiometric matrix of the sub-graph – and identifying the corresponding *balanced flows*.

Fig.4 shows the largest sub-graphs of the full generator for which the stoichiometry possesses left null eigenvectors. Only fertilization reactions appear, corresponding to conversions

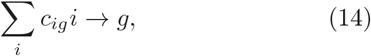

where *c_ig_* with 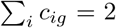, ∀*g* indicate the gametes that form each diploid. We denote a single event of each such fertilization as a basis vector for flows *ê_*i*_1*i*2→g__*, for and *i*_2_ the haplotypes indicated by *c_ig_*. The quantities conserved by fertilizations are allele counts. For random mating, these are

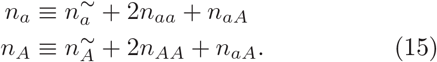

**FIG 4:**
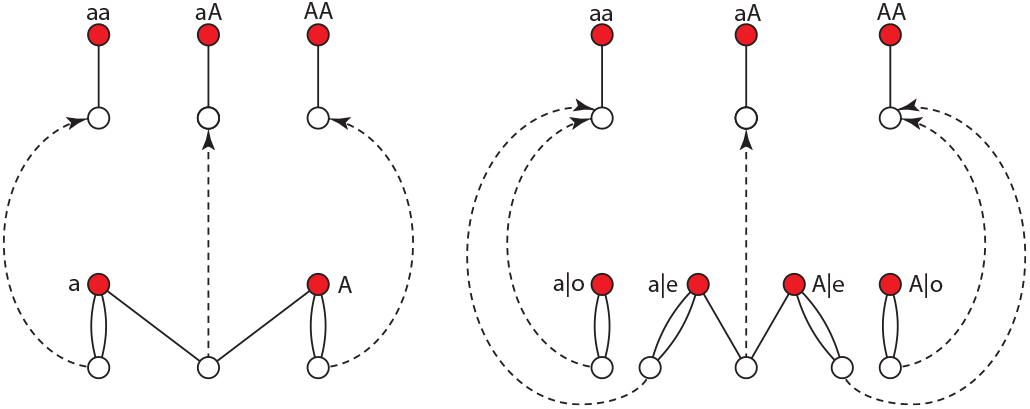
The sub-graphs from the generators of Fig. 2 that conserve allele counts. Note that only types *a* or *A* are con-served quantities in either graph, as *a* | o, *a* | *e, A* | *o*, and *A* | *e* are not allele types in the diploids.

For selfing, two separate sub-populations of haplotypes are conserved independently,

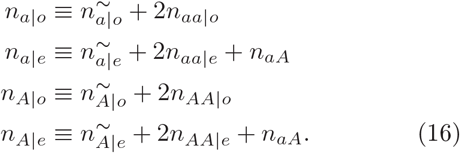

The allele-conserving fertilizations may be combined with the species-generating gametogenesis events to make autocatalytic flows completing the lifecycle, as shown in Fig. 5. In the figure, basis elements for the space of all flows are shown by integers attached to the reactions, indicating how many events of each reaction type make up a flow. Distinct basis flows are lettered with different colors.

**FIG 5:**
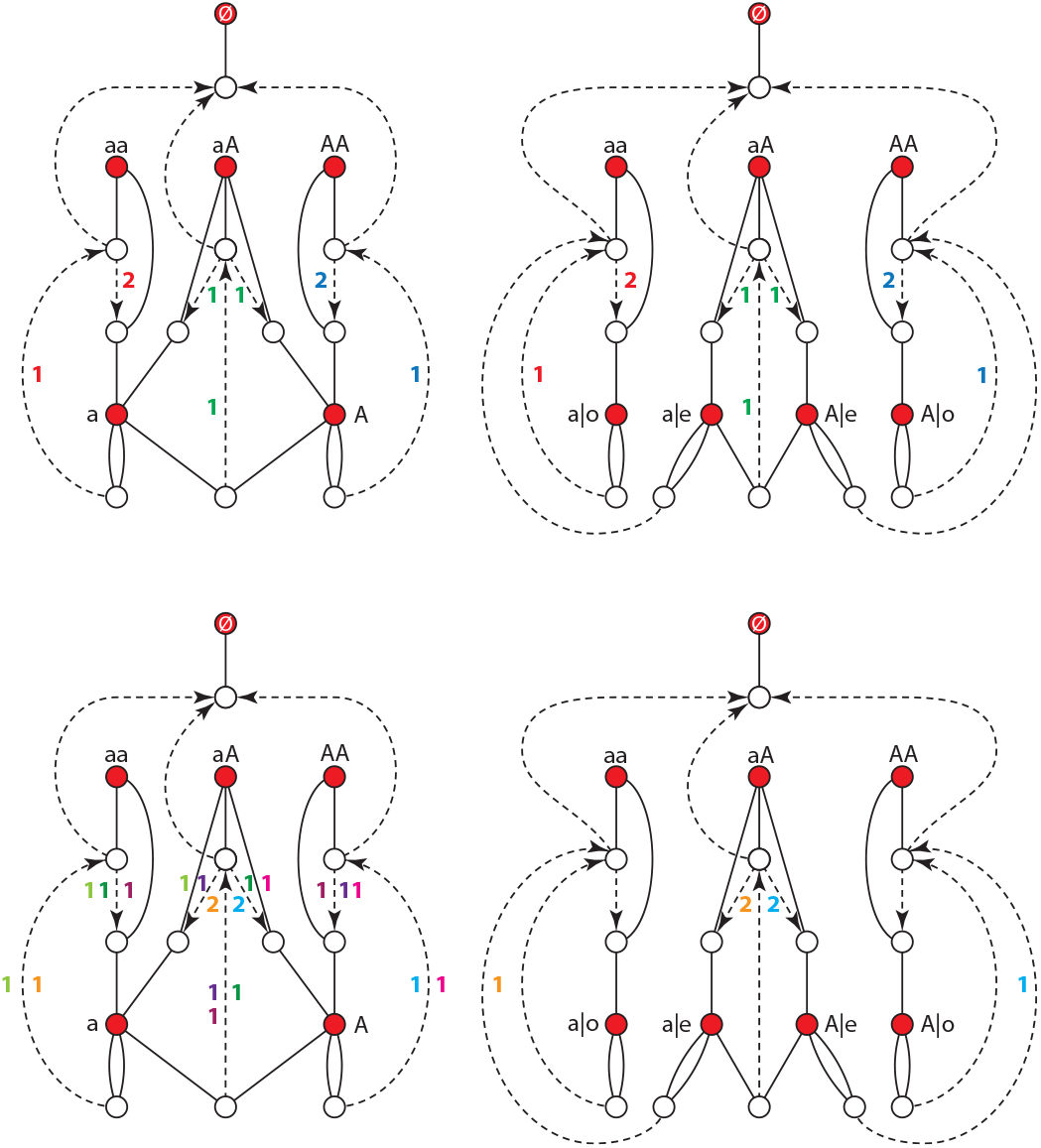
Autocatalytic flows, with allele counts conserved in the subgraphs of Fig. 4. Integers with the same color are the event counts in a single flow. The top pair of graphs show flows that are autocatalytic at the species level, for the three diploid genotypes *g*. The lower pair of graphs show additional flows that are cross-catalytic, depending on the presence of one species to produce another. The selfing and random-mating graphs share two purely cross-catalytic flows from the heterozygote to either homozygote. The random-mating graph has five additional flows that are mixed autocatalytic and cross-catalytic with respect to species.

All of these flows are autocatalytic in the network sense. They may be further partitioned into three subclasses: autocatalytic at the species level, if the parent species needed to execute a flow produce an offspring of the same type; purely cross-catalytic with respect to species if the offspring is different from any parent that contributed gametes to it; or mixed if one parent was the same type as the offspring and one different.

Letting *e_g→g_,i* denote single events of the gametogenesis reactions, the basis elements for autocatalytic flows may be written as linear combinations of elementary reactions. We introduce a superscript *G* ∈ {R, S}to refer to the graph for random mating or selfing, and write these basis vectors as 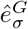, where *σ* is an integer (assigned arbitrarily) distinguishing flows. For random mating, the distinguishable autocatalytic flows are the combinations

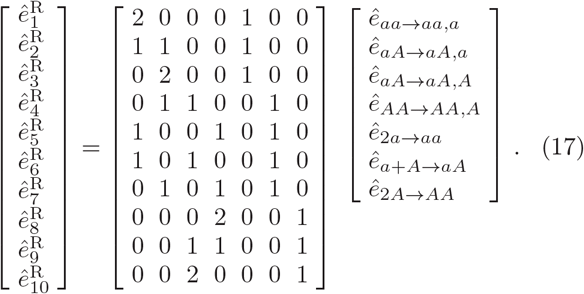

Any autocatalytic flow on the random mating graph can be written in the form 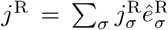. In place of *n_i_* of Eq. (15), which are conserved within the sub-graph of Fig. 4, the flows (17) conserve gamete counts 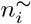, which we may hold at zero to leading order in *1/κ* to reflect the approximation (9) at large *κ*.

The ten flows in the basis (17) are not all independent. Five linear combinations of basis vectors exist:

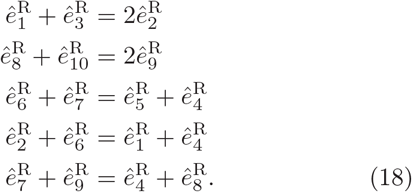

Seven linear combinations of currents in the graph are set by the rate equations (8) and (10)

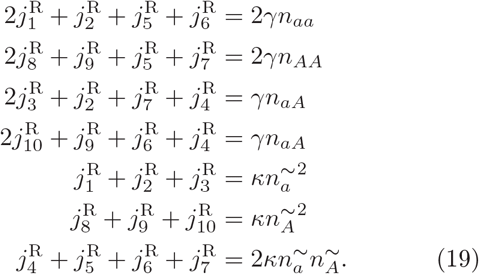

Two of these reflect the large- *κ* approximation 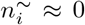 in Eq. (9), already incorporated in the linear combinations (17), and the remaining five determine the five linearly independent autocatalytic flows responsible for the rate equation (3) for random mating.

The linear dependencies (18) imply that stoichiometry alone cannot assign unique flow values to all basis elements in the set (17), reflecting the biological reality of reproduction without replication: no single path of a haplotype from a diploid parent, through the gamete phase, to the type of the offspring, is entailed in sexual reproduction. If such a unique path were entailed, fitness would be definable in Fisher’s apportionment sense for every entity in a lifecycle and further summary statistics would not be needed.

For the case of ephemeral gametes, however, in which 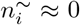, we may derive a refinement of the current constraints (19) by noting that for each flow 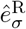, we may define an indicator function 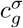 with 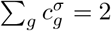, which is non-zero only for genotypes *g* that are ancestors in the flow *σ*. If we require, of each current 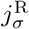, that

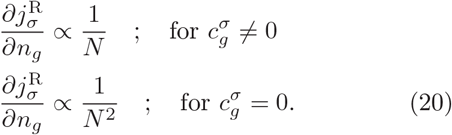

the resulting solutions reflect the catalytic dependency of each flow on the species number in the parent generation:

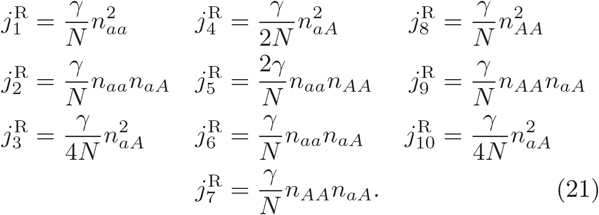

A similar and simpler construction can be carried out for the selfing graph in Fig.5. Gametogenesis occurs in more forms, but fewer overall flows arise as linear combinations of gametogenesis and fertilization,

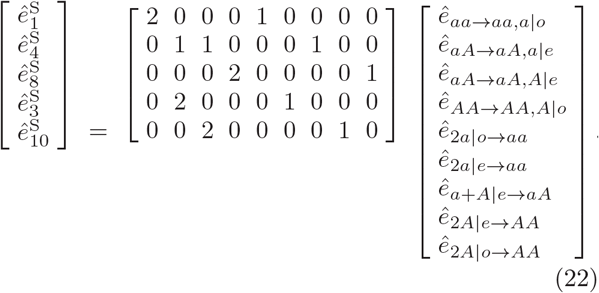

All five flows (22) are independent.

Seven currents are again assigned by (11) and (13),

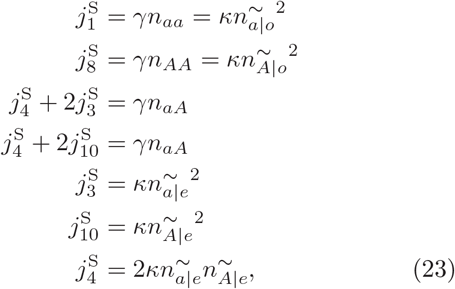

two of which are redundant with conservation of 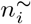 already provided by the basis elements (22). The resulting flow solutions, counterpart for selfing to the solutions (21) for random mating, are given by

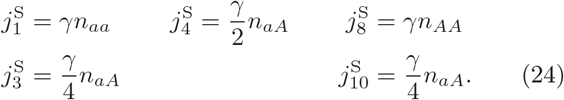

### B. Asymmetric gametogenesis: fitness at two stages in a lifecycle

The same procedure may be followed to analyze richer lifecycle models in which gametes carrying the two alleles *i* ∈{*a, A*}are produced at different rates. We will incrementally relax symmetries in the generators of Fig. 2, employing progressively more of the information-carrying capacity of the basis elements (17,22).

In the first refinement, gametes in either graph *G* are produced at unequal rates

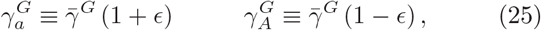

independent of whether the parent is homozygous or heterozygous. The resulting population process possesses a well-defined haplotype fitness, which nonetheless cannot be captured in additive fitness at the diploid phase of the lifecycle. We introduce a notation

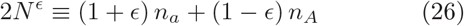

for a quantity that replaces 2*N* at many points in the analysis from the symmetric model.

The decomposition into currents corresponding to equations (21,24) for asymmetric gametogenesis is given in equations (B1,B2) in App. B1. The reproduction terms in the rate equations (3) from random mating are replaced by the three sums of autocatalytic flow currents

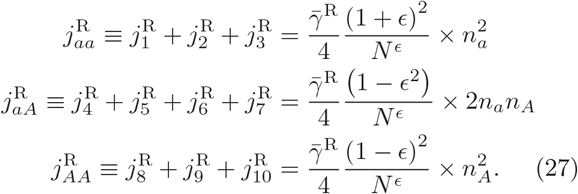

For selfing, the reproduction terms in the rate equations (4) are replaced by the sums

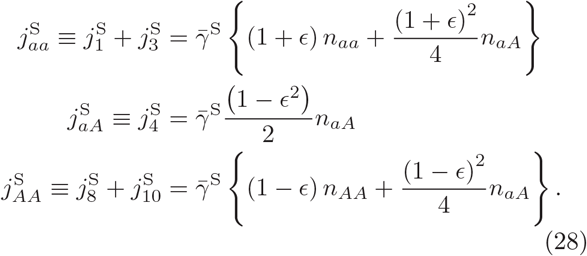

While the condition (20) makes it possible to identify ten distinct currents in Eq. (21), the redundancy of the coefficients of different monomials makes clear that all ten are not needed. As equations (27,28) show even for the more refined model, only six different monomials are actually required to span the flows of both graphs. We show next how to identify the combinations that are needed for a full covariance expansion in a given case.

### C. Bases and regressions

The autocatalytic currents in equations (21,24) or equations (27,28) may be written as regression coefficients times monomials, as

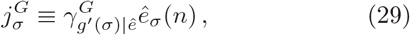

where *ê_σ_*(*n*) denote monomials in one or more of the following five count variables

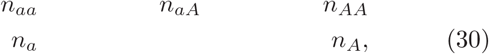

*g*′(*σ*) denotes the genotype produced by the current *σ*, and 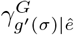 is the regression coefficient. The values of 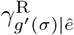 and 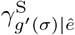 respectively from equations (27,28) are given by

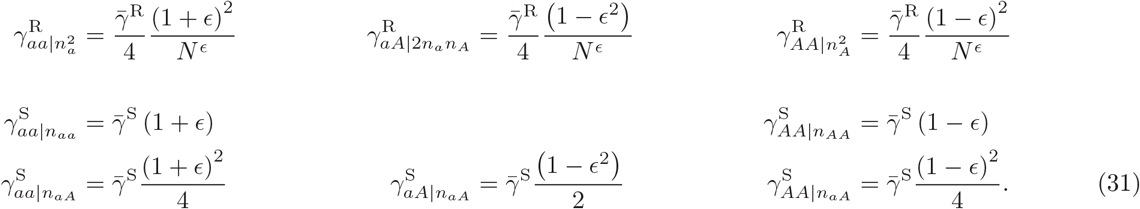

### D. Meiotic drive in heterozygotes

Finally we consider one further reduction of symmetry that gives a model for meiotic drive. In either graph, the rates of gametogenesis from homozygotes 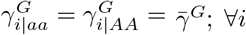; ∀*i*, but from heterozygotes allele *a* out-competes allele *A* in gametogenesis,

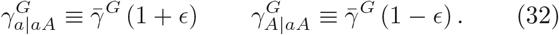

For this model, the fecundity contributing to fitness in Eq. (5) for all diploids is constant at 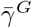, identifying the only relevant notion of competition as arising *within*the haploid lifecycle phase. The model of meiotic drive will be contrasted in Sec.VIID with a model of uniform asymmetric gametogenesis but non-uniform mortality to represent competition *between* levels.

The decomposition into currents corresponding to equations (21,24) for this model is given in equations (B3,B4) in App.B 2. The sums of currents corresponding to Eq. (27) for random mating now become

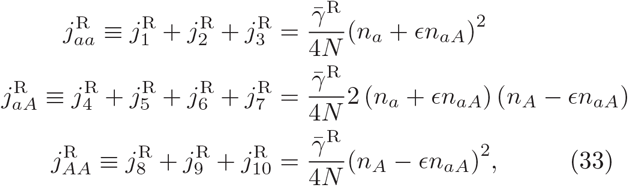

while those for selfing corresponding to Eq. (28) become

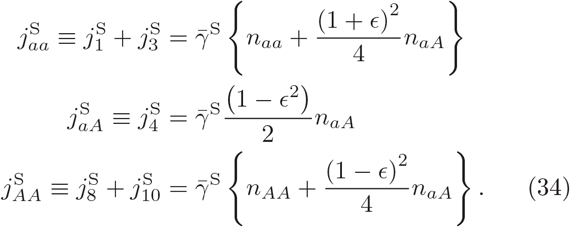

A common underlying form is preserved for each lifecycle; only the basis monomials and regression coefficients shift between cases.

## VI. THE PRICE EQUATION

A generalized Price equation can be constructed directly from the basis monomials and regression coefficients of the autocatalytic flows in Eq. (29). To compare that construction to the conventional construction over allele indices *i*, we first write down the Price equation with additive genic fitness, and then show, term by term, how a mis-specified regression on replicating genes, requiring an *ad hoc*“environment” correction, is replaced by an exact regression on flows that requires no such correction.

### A. Additive fitness in diploid reproduction models

To write down a conventional Price equation it is necessary to apportion offspring counts to parents according to the parents’ types. For a single-locus diploid sexual lifecycle this problem is simpler than it will be in cases with more general redistributions, because the fixed ploidy assigns a fitness of 1/2 to the parent per gamete that succeeds in fertilization. For uniform gametogenesis the fecundity rate accrues as in Eq. (5) over the lifespan.

To carry out the same apportionment more generally, one recognizes that each of the current assignments 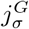 in Eq. (21) or Eq. (24) propagates some allele from a unique parent genotype *g* to an offspring genotype *g*′(*σ*). Summing these currents with coefficients 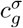 from Eq. (20) then defines an additive fecundity for type *g*. The structure of the apportionment is dictated solely by the stoichiometry of the flow; all dependence on population state comes through the regression coefficients 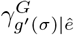.

The sums of currents, respectively of diploids into, or gametes out of, each genotype *g* introduce three new quantities:

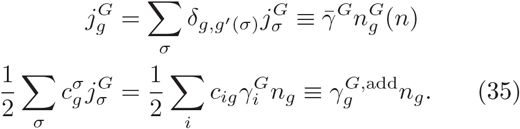

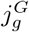 is the sum of all currents that produce genotype *g*. It defines a profile 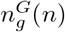 deposited at rate 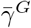 into the population. 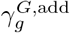 is the corresponding additive component of fecundity, generalizing Eq. (5) to non-uniform gametogenesis and generally introducing population state-dependence as a result.

### B. Characters and the Price decomposition of character change

Let *χ* ≡ (*χ_g_*) be a vector of characters defined on genotypes *g*. Then the mean additive fitness and change in *χ* accounted for by additive fitness are

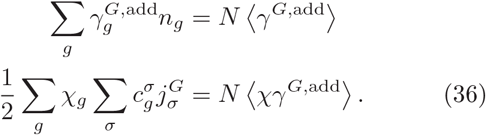

Now re-introduce mortality *μ* ≡ (*μ_g_*) from equations (3,4), with mean 〈μ〉. For generality, also allow for the possibility of partial self-incompatibility so that a superposition of the mating systems *G* ∈{R, S} may determine total population change. Then total number *N* evolves as 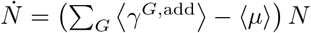.

Price’s *character change* Δ_χ_ can be defined strictly from stoichiometry for each flow *σ* on a graph *G* as

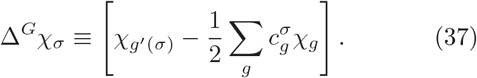

In terms of the quantities (35) and (37), the rate of change of〈χ〉 is then written in the canonical Price equation as

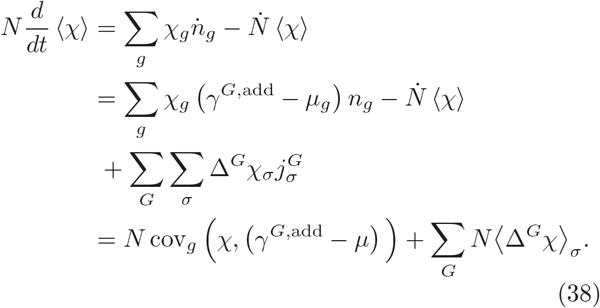

The first term in the final line of Eq. (38) is the covariance term in Fisher’s fundamental theorem, and the second term is variously named the “environment”[17] as well as the “character change” term.

### C. From character change to a covariance on flows

As is well understood [14], an additive fitness model is not dynamically sufficient [34] to explain population trajectories under a process with underlying gametic fitness if the lifecycle also contains a redistributive stage such as fertilization. We now show how the mis-specification is overcome by using a basis of autocatalytic flows.

The average of Δ^G^_*χ*_ from Eq. (37) is a difference, for each *χ_g_*, of the flows into and out of genotype *g*. By Eq. (35), the term proportional to the constant part of *χ* can be written two ways,

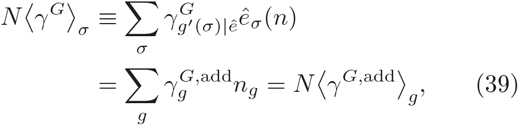

in the flow index σ or as an average over genotypes *g*. Adding and subtracting Eq. (39) from *N* Δ^G^_*χ*_❭ gives

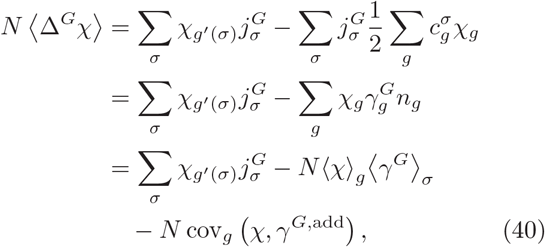

recovering on the input side the additive covariance in Eq. (38). The total change of ❬χ❭ from fecundity is the combination present originally, if additive fitness had never been separated,

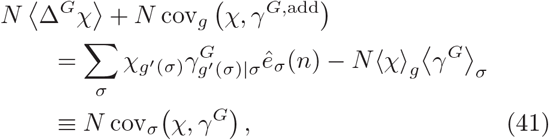

seen to be simply a covariance on index *σ*.

In other words, the Price equation (38) with additive fitness and character change re-arranges to a pure covariance expression at two levels:

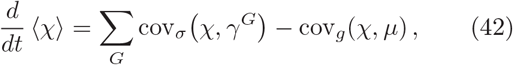

in index *σ* for fecundity and in index *g* for mortality (where *g* is also the index for the basis of independent flows to the null object Ø). **Eq. (42) is the first main result of the paper.**

### D. Current basis for covariance in the notation of the examples

In the examples, with three dynamical components, a symmetry-based coordinate system separates population size, additive fitness, and dominance effects. Let *T* stand for a trace component of any character χ, and || and ⊥ for the components that are respectively antisymmetric and symmetric under *a* ↔ *A*. Then *Ṅ* and *d* ❬χ❭ /*dt* can be written

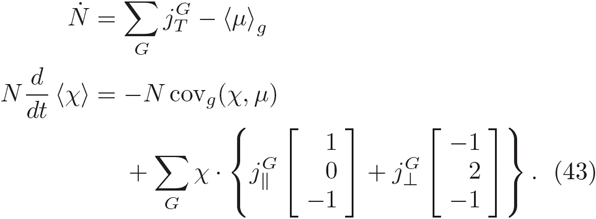

In the basis set of Eq. (17) for random mating, the three components are sums

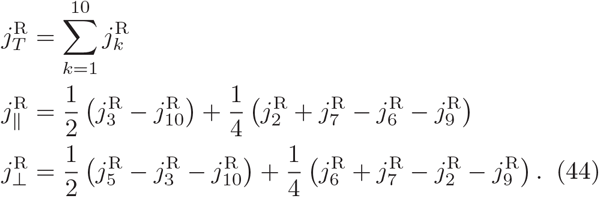

In the basis set of Eq. (22) for selfing, they are

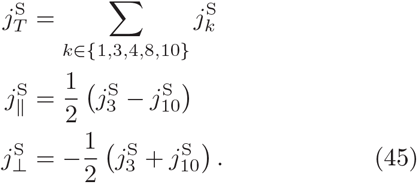

## VII. COMPARISON OF PRICE EQUATION FORMS IN DIFFERENT SELECTION REGIMES

App.A reviews properties of additive regressions against the differential mortality component of fitness among diploids. In this section we consider the two forms of asymmetric gametogenesis introduced in Sec.V. Uniform asymmetry leads to well-defined allelic fitness that is mis-specified in diploid additive models but handled correctly in Eq. (42). Meiotic drive makes no contribution to the additive fitness, and occurs entirely through the constructed character of heterozygosity. Framed in terms of multi-level selection, the two cases contrast selection acting between levels versus acting within a level.

### A. Additive diploid fitness approximating genic selection

Uniform asymmetric gametogenesis rates (25) lead, as expected, through Eq. (35) to additive fitness contributions

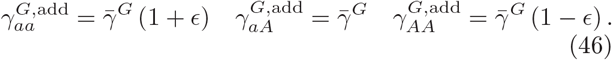

Fisher’s covariance term in Eq. (38) evaluates to

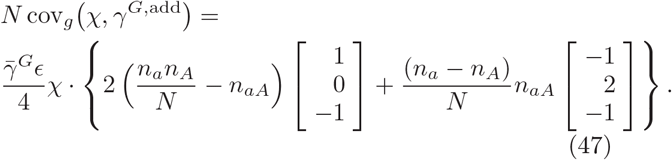

The analysis of the additive approximation for gametogenesis mirrors exactly the analysis in App. A for mortality. The “allelic components” of fitness 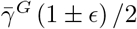 accounting for the diploid coefficients (46) are recovered (up to a common offset) in the components (*α_a_, α_A_*) of an additive regression in what is called the “*α*-representation” [14]. (The difference of the allelic components 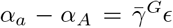 is recovered from the counterpart to the value *λ* in Eq. (A12).) The actual trajectory components *d* log *n_i_*,/*dt* from the first summand on the right-hand side of Eq. (47) are generally related to the regression coefficients (*α_a_,α_A_*), by state-dependent factors derived in the appendix.

In particular, the functional dependence (*n_a_n_A_*/*N* – *n_aA_*) implies state-dependent values of *d* log *n_i_*/*dt*, obscuring the property of actual trajectories that these derivatives are the constant gametic fitnesses 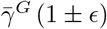. The “character-change” component 〈Δ^*G*^_*χ*_〉, needed to reveal the allelic nature of fitness, is in this case better described as absorbing the mis-specification of the additive diploid model (along with other properties of the mating system). For random mating, the full Price equation (38) takes the form

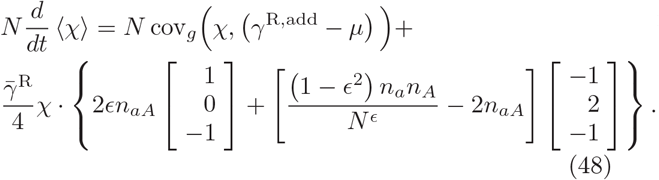

For selfing, the form is

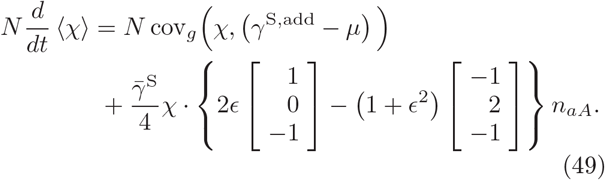

The coefficient in the first summand, 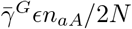, is the same in either mating system and cancels the spurious factor of *n_aA_* in the first summand of the additive covariance (47).

Note that any population-genetic model also defines a quantitative model if we take indicator functions for genes or genotypes as the quantity characterizing the phenotype, upon which Robertson’s second theorem follows from the Price equation [11]. Doing so for gametic selection we would identify 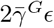 as the gametic fitness difference, and *n_aA_*/4*N* as a coefficient of heritability, inevitably state-dependent because it refers to the likelihood of heterozygosity which is not attributable to gametes.

### B. For asymmetric gametogenesis a multilevel covariance is directly recovered from the hypergraph Price equation

The actual genic fitness values are not the “allelic components” of the additive diploid fitnesses (46), but those obtained from the current decomposition in either of Eq. (27) or (28), which are larger by a factor of 2:

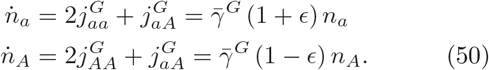

In a regression model at the gene level, simply ignoring the existence of the diploid phase in the lifecycle, the contribution to 〈χ〉 from the projection onto alleles can be written

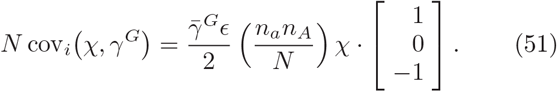

The expression (51) gives no information about the components of *χ* that distinguish homo-from heterozygosity, and thus the effect of the mating system on this component. However, if subtracted from cov_*σ*_(*χ, γ^G^*) in the Price equation (42), it leaves a remainder term only describing the effects of the mating system. For random mating,

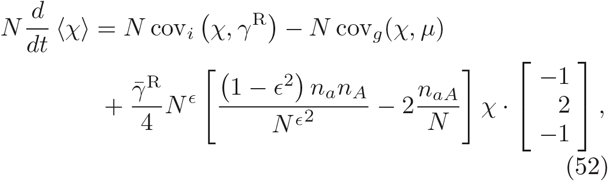

while for selfing

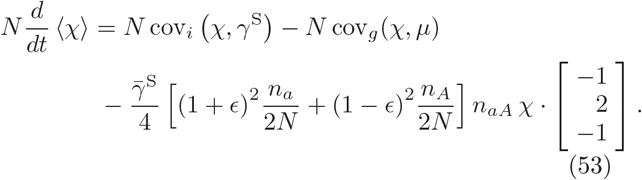

Both expressions (52,53) include multilevel covariance relations, with covariance on the allele index *i* for fecundity, in which genic fitness is exact, and covariance on the diploid genotype *g* for mortality, which is linear in *n_g_*.

The different mating systems attract onto the invariant manifolds respectively for Hardy-Weinberg equilibrium or for homozygosity, described in App.A 1. To the extent that they express purely redistributive effects of the reproductive cycle, they can be properly regarded as character-change terms in Price’s [1] sense, and the symmetric character modulated is dominance. The contrasting example of meiotic drive, however, considered next, shows the sense in which these terms are as naturally regarded as covariance expressions through Eq. (42).

### C. For meiotic drive there is neither diploid nor haploid reference fitness

For the gametogenesis rates (32) defining meiotic drive, we focus on the random-mating case. For selfing, meiotic drive is distinguished from uniform genic selection only by the presence or absence of an additive fecundity component for the diploid, which will be considered jointly with mortality in Sec.VII D.

For random mating, the generalized Price equation (42) is written directly as

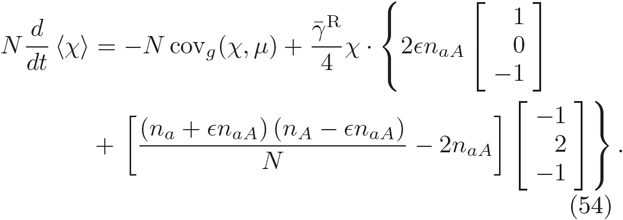

Eq. (54) again describes attraction onto a position in the Hardy-Weinberg equilibrium manifold, now at allele frequencies displaced α ϵ from the starting population *n*.

### D. “Competition” within and between “levels”

Following observations made in Sec.VD, let meiotic drive (32), with uniform mortality 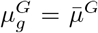, ∀*g*, define “within-level” competition between alleles *a* and *A*. A contrasting model of “between-level” competition^10^ can be obtained by combining the uniform asymmetric gametogenesis (25) with non-uniform mortality of the form

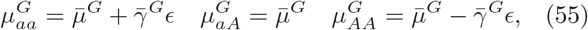

so that 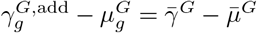; ∀*g*.

Additive fitness for diploids will be uniform in both cases, but for different reasons. Within levels, gametes may be asymmetrically antagonistic or may compete with each other for parental resources, leading to a stalemate in homozygotes and differential gametogenesis in heterozygotes. Between levels, one haplotype achieves faster production than the other in all contexts, but does so at the expense of the parent’s longevity, canceling its gains.

Under self-fertilization, competition within and between levels is indistinguishable. For either, the current components in Eq. (43) take the forms

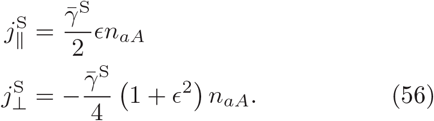

For random mating the two forms of competition are distinct. Under meiotic drive, Eq. (54) leads to currents

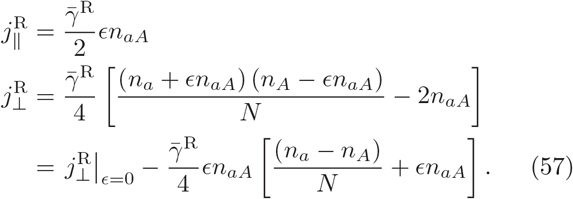

Eq. (57) may be compared against the exact form (48) for between-level competition. 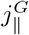 is the same for both cases and for Eq. (56). To make clearer the relation between 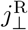 for within- and between-level competition, Eq. (48) expands to 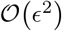 to give

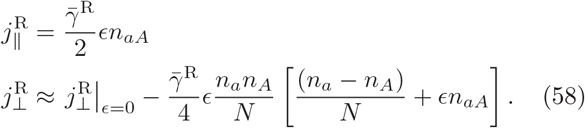

Fig. 6 illustrates the different consequences of the two forms of competition for population trajectories. A population of *AA* is invaded by haplotype *a* through the heterozygote. The new allele is first shown in the figure when genotype *aA* reaches 1% frequency. For relative allele fitness *ϵ* = 0.2 it takes 45 generations^11^ for haplotype *a* to sweep to ≈97.5%and 97.2%of the population, respectively for within- and between-level competition. The two alleles pass through equal frequency at respectively ≈26.6 and 26.85 generations. More visible in the figure, both populations sweep out trajectories on which the heterozygote is depressed relative to Hardy-Weinberg equilibrium, with between-level depression ≈4.25×as large as the within-level depression.

**FIG. 6:**
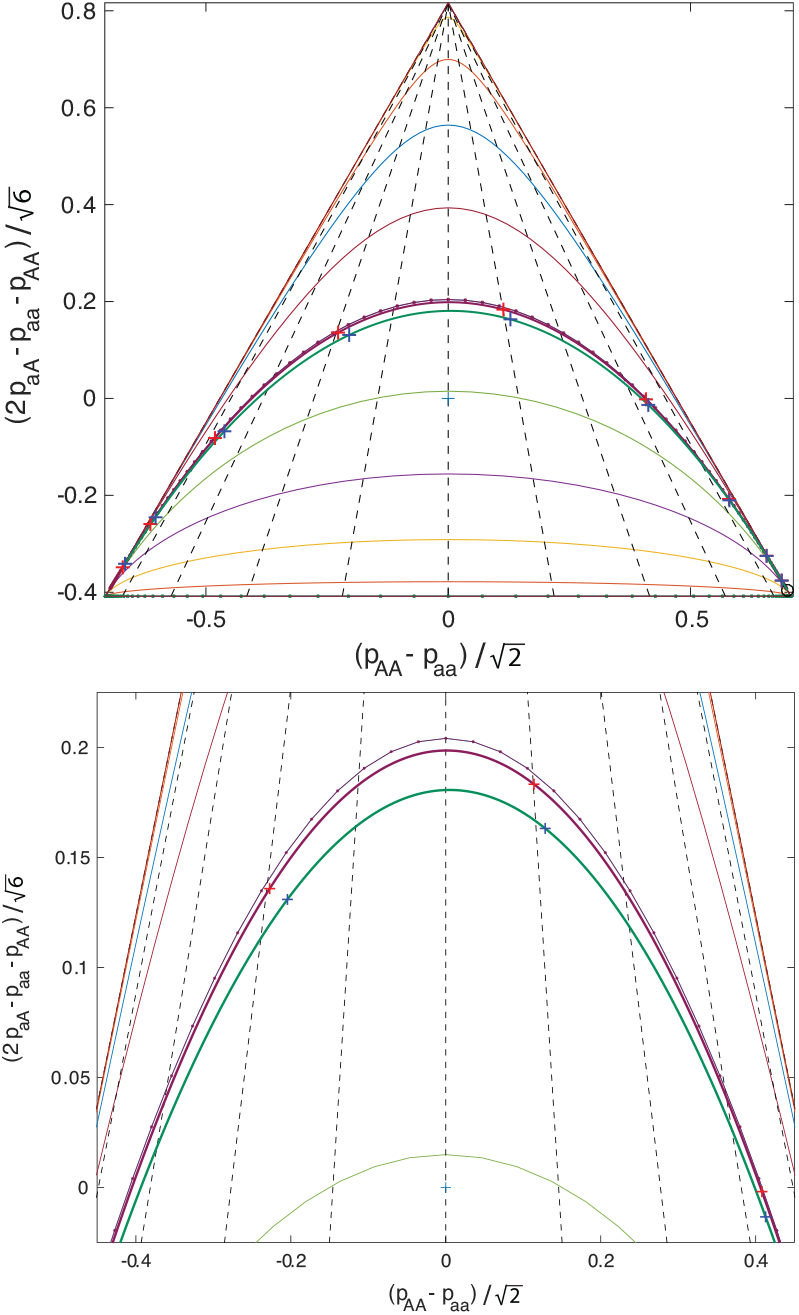
Comparison of within and between selection, for ∈ = 0.2. A nearly homogeneous *AA* population is invaded by a haplotype *a* in a heterozygote, which then sweeps to fixation. Heterozygotes are depressed relative to Hardy-Weinberg equilibrium, but the depression is about ≈4.25×as large for between-level selection as for within-level selection.

#### Remark

The meiotic drive model was constructed to have strictly zero differential fitness for genotypes *g*, and only a conditionally-heritable fitness differential for haplotypes *i*. Any effect of allele asymmetry on the population trajectory arises entirely through the constructed property of heterozygosis. Clearly there is no reason *not*to regard such effects as products of selection, yet the Price equation (38) referenced to the diploid phase of the lifecycle consigns these to the 〈Δ^G^_χ_〉 term. Expanding the inventory of regression coefficients to include constructive autocatalytic flows incorporates such effects naturally within the covariance of Eq. (42). Once the coefficients 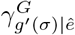 have been recognized as the same kinds of regression coefficients as fitness, simply on an expanded basis, not only selective effects in between-level as well as within-level competition, but the redistribution of the mating system itself, are channels of selection to be estimated by population-genetic methods through their contributions to population trajectories.

## VIII. A MEASURE FOR THE INFORMATION IN A POPULATION TRAJECTORY

Selection as a concept should refer to the aggregate effects of all the processes that put information into a population state through repetition of lifecycles of the members. Because the exact population state fluctuates, information should refer not to the exact state, but to a distribution in which the observed samples of states would be typical. A notion of added information further requires a reference distribution or null model, against which the evolved distribution is to be compared.

A measure that fulfills all three criteria is the *large-deviation function* [37] for an observed sample property from the null or reference distribution. The simplest example – not the one ultimately to be used here, but illustrative of the above criteria – might be the probability to sample a population with composition *n*′ of size 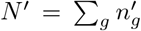 from a distribution expected to produce populations with some typical composition 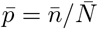 where 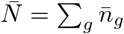. To simplify the example suppose that the sample size *N*′ is chosen externally, and it is only deviations in sample composition (the feature governing 〈χ〉) for which we seek an information measure.

The probability to draw a particular sample *n*′ by independent draws of members in any order is

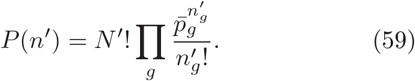

For distributions on discrete states such as populations, it can be shown [39] that the leading *N*′-dependence of −log *P*(*n*′) is the large-deviation function (LDF) for the sample *n*′. Using Stirling’s approximation for factorials,

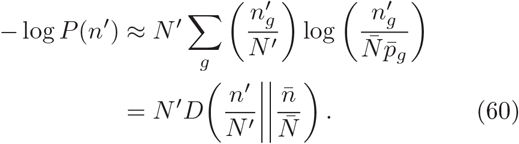

A defining feature of the LDF (60) is its separation into a factor *N*′ characterizing the *scale* of the sample, and a factor 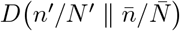 characterizing the *structure* of the deviation [38].

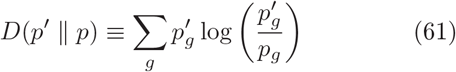

is called the *Kullback-Leibler divergence* of distribution *p*′ from distribution *p*. It is the negative of the relative entropy^12^ of p′ referenced to measure *p*, is positive-semidefinite, and vanishes only when *p*′ = *p* [66]. The LDF (60) extends the well-understood interpretation of the log-improbability of a sample as a measure of information [67, 68], to a measure of the divergence of a tilted distribution in which *n*′ would be a typical sample, from a reference distribution in which *n* was typical.

Here we will not want the large-deviation function of a population *state*from some reference state, but rather the large-deviation function for an entire population *trajectory* from the trajectories typical under some null model. There are two main reasons a trajectory entropy is the desired route to a measure for the information we introduced as being “in a population state”:

1. Selection acts through the *rate of change* of a population state, and it is this rate of change – rather than the accomplished change over some arbitrary time interval – from which we wish to compute an information-carrying current. Attempting to use a measure such as (60), perhaps with *n* being the present state and *n′* = *n*+*dtṅ* the “informed” state, gives a divergence that does not compose over time regularly in *dt*.
2. The foregoing examples show how a population with only a few types, but even a modestly-structured lifecycle, may evolve under a variety of different selective regimes. A regression model of *ṅ* at any one time is inherently underdetermined: a sub-manifold of generating parameters can be assigned at any single population state. This under-determination of a model from *ṅ* is an important reason to minimize the number of regression coefficients, which projection onto additive models does, even at the cost of mis-specification. In contrast to a single tangent vector *ṅ*, an extended population trajectory is formally infinite-dimensional, thus sufficient to identify any finite-dimensional generating model under sufficient resolution and sample size.

Many methods exist to study ensembles over trajectories and to compute the trajectory LDF. Here we use a method derived from the Liouville operator that evolves the moment-generating functions of time-dependent probability distributions over states. The method is variously known as the *momentum-space WKB approximation* [69], the *Hamiltonian-Jacobi method* [70], or in a diffusion approximation as *Freidlin-Wentzell theory* [71]. It may also be derived from a variety of 2-field functional-integral methods [72–76], reviewed in [39, 77, 78]. Its application to population processes is introduced didactically in [40, 55], so only a brief review sufficient to define terms and major constructive steps is provided here in App.C.

### A. The 2-field functional integral construction of generating functions and functionals

The core object in the construction is a time-dependent probability distribution *ρ_n_* on population states, evolving under a master equation^13^

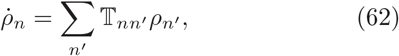

with transition matrix 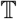. A moment-generating function (MGF) Ψ(*z*), for *z* ≡ (*z_g_*) a vector of complex coefficients, is defined as

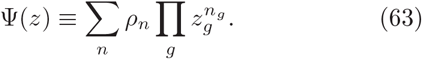

The master equation (62) implies that Ψevolves under a *Liouville equation*

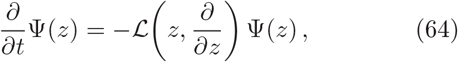

in which the Liouville operator 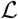 is constructed from the elements of the transition matrix 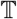. By taking the quadrature of Eq. (64), Ψ(*z*) at any time *t* = *T* > 0 can be computed from initial data Ψ_0_(*z*) at *t* = 0.

By a standard derivation sketched in App.C, the MGF can be written as a 2-field functional integral of the form

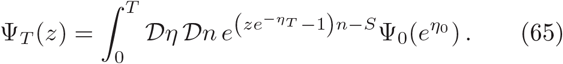

In Eq. (65), *n* ≡ (*n_g_*) and *η* ≡ (*η_g_*) are vector-valued fields of integration on the interval [0, *T*], 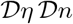 is a functional measure on *η* and *n*, *e*^±*η*^ is the component-wise exponential, and 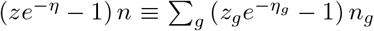 is the vector inner product of (*z* – *e^η^*) regarded as a row vector and *e^-η^n* regarded as a column vector. In the following, where vector-valued fields are written in products, exponentials, or logarithms, these are to be understood as component-wise products, exponentials, or logarithms. Subscripts *η_T_* or *η*_0_ in Eq. (65) indicate the values of the field *η* at the limiting times *t* = *T* and *t* = 0.

The form of the functional integral (65) is general for population processes on a given state space. All information from the transition matrix 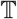 for a particular process is expressed in the weight functional *S*, which has the form of a Lagrange-Hamilton action functional, with 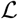 as its Hamiltonian:

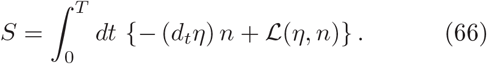

The classical function of fields 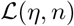 in Eq. (66) is obtained from the Liouville operator 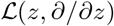 of Eq. (64) by the argument substitution *z_g_* → *e^η_g_^, ∂/∂z_g_* → *e*^−*η_g_*^*n_g_*.

#### 1. Cumulant-generating function and generating functional

We will refer to *z^n^ρ* in Eq. (63) as a *tilt* of the distribution *ρ*.^14^ Denote by *θ* = log *z* the component-wise logarithm. Then the cumulant-generating function (CGF) *ψ*(*θ*), with natural argument *θ*, is the log of the MGF:

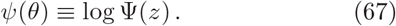

The expectation of the state *n* in the (normalized) tilted distribution *z^n^ρ_n_*/Ψ(*z*) is the gradient of the CGF,

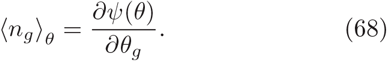

If *ψ*(*θ*) is convex, the function (68) is invertible to a function *θ*(〈*n*〉). The Legendre transform

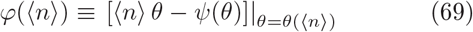

is the large-deviation function for distribution *ρ*, providing the formal derivation behind the intuitive example (60). By construction its gradient gives

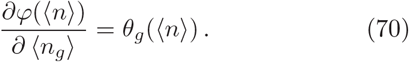

It follows that the minimizer of *φ*(〈*n*〉) is the expectation under the original distribution *ρ*.

The CGF (67) of a single argument *θ* at a final time *T* may be generalized to a cumulant-generating functional on trajectories. Let *σ* ≡ (*σ_g_*), termed a *source*, be a vector-valued real function on the interval *t* ∈ [0, *T*]. Then the functional CGF *ψ*[*σ*] is defined from a variant on the 2-field integral (65) as

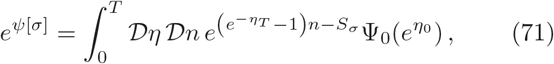

in which the source-perturbed action

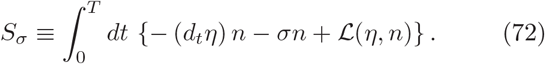

The source term *σ* in Eq. (72) tilts the extended-time distribution over trajectories, just as an endpoint parameter *θ* can tilt a single-time distribution. Since it is possible for the history *σ* to include *δ*-function contributions at any time, we omit the explicit surface argument *θ* of Eq. (67), and set *z* appearing in Eq. (65) to value *1^T^* in Eq. (71).

The trajectory generalization of the gradient relation (68) is the variational relation under a variation *δσ*:

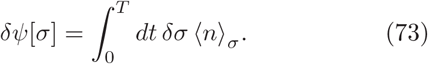

Like the single-time CGF, *ψ*[*σ*] possesses a functional Legendre transform

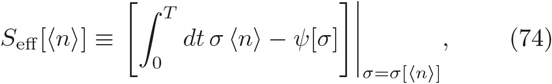

which is the large-deviation function for trajectories in a time-dependent distribution evolving under the transition matrix 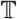 of Eq. (62). *S*_eff_[〈*n*〉], a functional of an entire averaged history 〈*n*〉, is called the *stochastic effective action* [39] (the reason for this designation will become clear shortly), and possesses the dual variation under variation *δ*〈*n*〉:

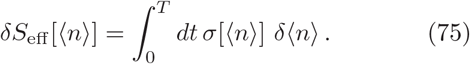

The minimum of *S*_eff_ identifies the actual mean trajectory 〈*n*〉 in the original distribution evolving under 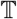.

#### 2. Saddle-point approximation of the large-deviation function

The functional CGF and LDF incorporate in principle all orders of correlated fluctuation in the population state, not only contemporaneous but also with time-lagged correlations. However, these are laborious to compute in general, and the leading exponential dependence of both the CGF and LDF can often be adequately approximated by saddle-point methods that are much simpler and still deceptively powerful.

The saddle-point approximation to a functional integral such as Eq. (71) is given by evaluating the integrand at a stationary path 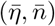 of *S_σ_* solving the Hamiltonian equations of motion

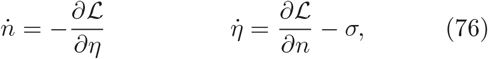

along with boundary conditions 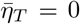 = 0 (for the choice of surface argument *z* → 1*^T^* in Eq. (71)) and a value for 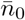 determined by Ψ_0_ and the solution value for 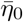. The saddle-point approximation to the mean is 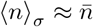. Using the boundary conditions to subtract a total derivative *d_t_* (*ηn*) from Eq. (72), the saddle-point approximation to the stochastic effective action becomes

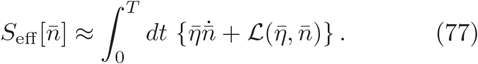

Although the explicit *σ*-dependence of Eq. (72) has been canceled in the Legendre transform (74), both 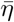 and 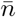 depend functionally on *σ* as solutions to Eq. (76).

Preservation of the norm 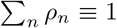 under stochastic evolution (62) is equivalent to the condition Ψ(1) ≡ 1 for any initial data Ψ_0_. This condition translates in the saddle-point approximation to the property of all Liouville functions for stochastic processes that 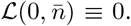. The manifold of solutions (76) at *σ* ≡ 0 with 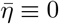 comprises all solutions to the deterministic rate equations for the population process.

#### 3. Quadratic expansion of the effective action in small tilts

The LDF (77) applies to any degree of deviation of n from the *σ* ≡ 0 background, but the nonlinear rate equations (76) and functional Legendre inversion (74) are generally nonlocal in time and may be difficult to compute. For small *σ*, however, the expansion of the LDF to 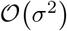 is an integral of a time-local function.

Denote the background solution under 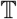 at *σ* ≡ 0 by *<u>n</u>*. Then the stationary-path equations (76) expand to linear order as

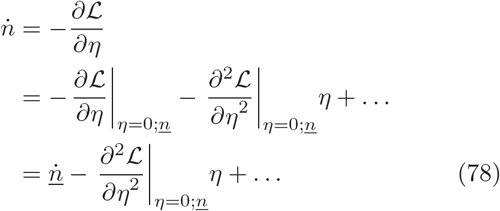

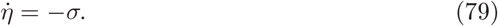

The function 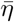 required to produce any deviation 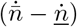 is computed from Eq. (78) by inversion of 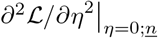, and the required source σ is then given by (79). Expanding the LDF (77) to second order in small *η* and 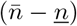 gives the approximation

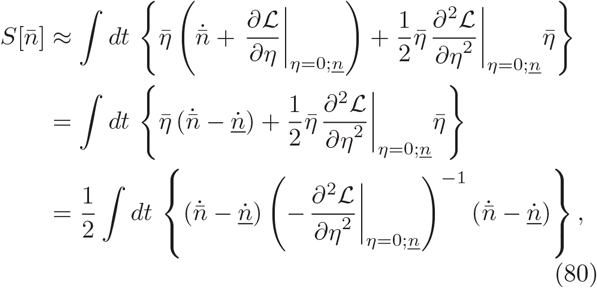

first due to Onsager and Machlup [82]. **Eq. (77), and its small-rate form (80) are the second main result of the paper.** The next two subsections will draw connections of this information measure to the conventional approach to evolutionary information through Fisher’s fundamental theorem, and provide explicit forms for the lifecycle graphs in the examples.

### B. Form of the Liouville function for lifecycles

To represent the graphical models of Fig.1 in functional integrals, two sets of paired fields must be introduced for the two sets of counting variables that define a state: {*η*_g_, *n_g_*} for diploid genotypes, and 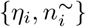 for gametes.

The simplest stochastic processes that will generate rate equations of the form in Sec.IV B (and the generalizations in Sec.VB and Sec.VD) are those equivalent to chemical master equations, in which each event of death, gametogenesis, or fertilization occurs independently.^15^ In addition to the coefficients *c_ig_* introduced in Eq. (14), introduce a second set of coefficients *f_g^i^_*_1^*i*^2_ equaling 1 for *i*_1_ and *i*_2_ the two haplotypes in *g* in an arbitrary (but fixed) order. These coefficients along with mortality rates *μg* and gametogenesis rates *γi_g_* (to gamete *i* from diploid *g*), are defined relative to each graph *G*. The Liouville function for the minimal process is then

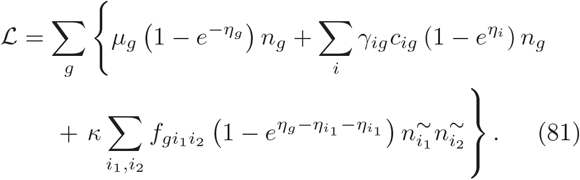

In solving the rate equations in Sec.IV B *et seq*., it was possible to eliminate the explicit variables 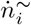 which remain near zero. A similar elimination can be carried out in the functional integral (71) by first integrating out the conjugate fields *η_i_* to produce approximate functional *δ*-functions which then set the field variables 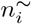 to the functions 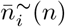 (*n*) from Eq. (10) or its counterparts for other cases. The net effect is to replace Eq. (81) by a generator with only fields {*η_g_*, *n_g_*} as arguments, of the form

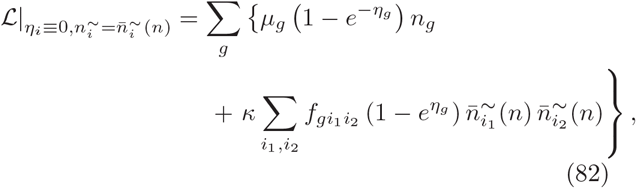

which then serves as the starting form for the saddle point approximations (77) and (80).

The first partial derivative 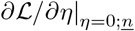 from Eq. (82) recovers the rate equations (3, 4) or their generalizations for 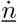. The second partial derivative required to evaluate Eq. (80) takes the form

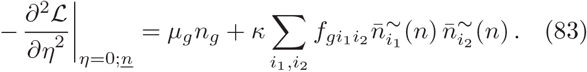

The matrix

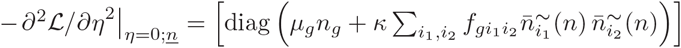

is the covariance of a Langevin field in the stochastic differential equation, representing the sampling noise rate of the underlying population process.

The information measure (80) derived from generator (81) is useful when 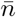 is an observed trajectory, and *<u>n</u>* a trajectory produced by a model with which the actual generating process may be confounded at an initial time. The divergence of the two trajectories over time measures the rate of accumulation of disambiguating information with repeated observations.

### C. Information referenced to the martingale

If we take the point of view expressed in the extended Price equation (42) – that the redistributing steps in the lifecycle are channels for selection as much as death or gametogenesis are, and similarly eligible to be inferred from trajectory statistics – the relevant null process against which to measure information input is a neutral process formalizing the notion of minimal prior assumption.

A neutral process for a population about which nothing is known except its current state is a martingale [83], in which the expected future state equals the current state. A neutral process formulated in terms of lifecycles might model each type of observed organism as a replicator, corresponding to the graph components in Fig.3, augmented by reactions *g* → Ø representing death of each diploid. The Liouville function for a birth-death process [79] with only replication and mortality is

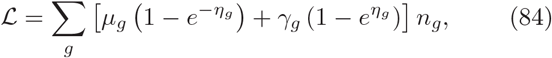

with rate equations and sampling variance

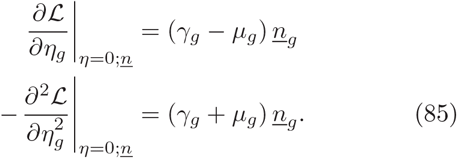

If the birth-death model treats types symmetrically, 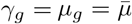, ∀*_g_*. The quadratic LDF (80) about the constant background *<u>n</u>* is then

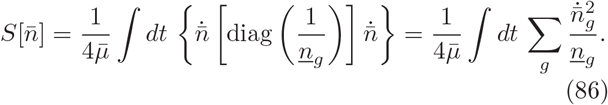

Consider the initial-time problem when the null model is set as 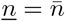 of the observed trajectory, and we wish to assign a rate of information incorporation to the tangent vector 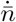. For a simple replicator with lifetime 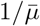, and fitness identified (as in Equations (6,7)) with 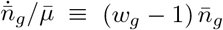, the initial integrand in Eq. (86) becomes

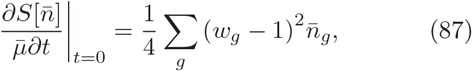

equal up to a constant factor to Fisher’s variance of fitness, in which the matrix 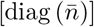 is the Fisher metric [84], also called the *Fisher information*.

**Remark:** The Onsager-Machlup information rate (87) coincides with the Fisher information because both have a common construction. The inverse of the Fisher metric, [diag(1/*n_g_*)] is the Hessian of the *f*-Divergence 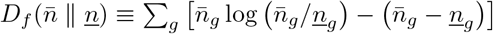 on either argument) at 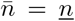 = *<u>n</u>*. The *f*-divergence is in turn the Kullback-Leibler divergence of a product of Poisson distributions with parameters 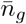 from one with parameters *<u>n</u>_g_* (see [55] for this derivation and more context). As developed by Amari and Nagaoka [84], the Fisher metric is the Hessian of the CGF for such a distribution in coordinates *θ* and its inverse is the Hessian of the Legendredual LDF in coordinates 〈*n*〉.

The trajectory LDF (80) accumulates the divergence rates of two such distributions with tangent vectors 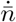 and *<u>ṅ</u>*. In setting 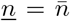, we have created a neutral process in which the covariance matrix for sample fluctuations within a generation can be checked to be [diag(*n_g_*)] /4 in the Gaussian approximation for fluctuations that defines the Onsager-Machlup action [82]. The covariance with additive fitness as used by Fisher also treats the population counts 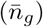 as sample weights during passage through one generation of a population’s lifecycle, and additive fitness is a regression coefficient matched to the population trajectory by a criterion of least squared error (see Sec.VII A). We may thus take the general information function (77) to define both the large-deviation context, and the correct formula for temporal accumulation, to derive variance of fitness as an information measure and to justify the squared error within a generation as a leading time-local expansion in the divergence of tangent vectors to observed and expected trajectories.

### D. Information channels

The trajectories 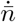 in Eq. (86) result from the autocatalytic flows 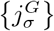 contributing to equations (44, 45) along with mortality rates 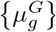. The variation in trajectory information resulting from variation of any of these regression coefficients is

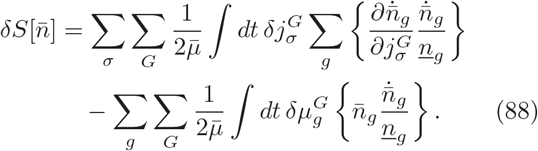

A marginal rate of information input at time *t*, through each reproductive channel *σ* or mortality channel *g* for a given lifecycle *G* is defined from the corresponding variational derivatives of 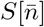 with respect to the regression coefficients

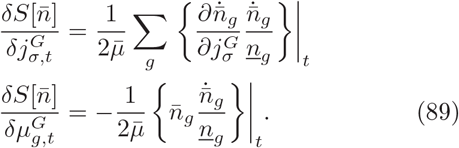

Whether increasing a given parameter imparts or dissipates information in a trajectory depends on whether the directional derivative along that parameter has positive or negative inner product (in the Fisher metric at *<u>n</u>*) with the tangent vector 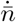 produced in aggregate.

The concept of information channel defined in Eq. (89) formalizes the interpretation of the Price equation (42) as a measure of the change in a population *due to* selection, including the fitness of replicators and also differentiable constructive operations within the lifecycle on an equal footing. The construction from the variational derivative applies equally to the general form (77), about arbitrary backgrounds (82), though the consequences of perturbations will generally propagate non-locally across time.

## IX. USE OF TRAJECTORY INFORMATION TO DISCRIMINATE SELECTIVE REGIMES

### A. Models under-determined initially become distinct through time

If two models are confounded in an initial observation because the regression of possible generating parameters against a single tangent vector 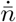 is under-determined, the maximum rate at which repeated observations of the population state can supply information to disambiguate them is measured by using one model as the generating process (82) for <u>*n*</u> and computing the information in the divergent trajectory 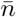 produced in the alternative model.

This situation arises even for the simple pair of models (3,4). In addition to the inherent confound of net fecundity and net mortality, if self-incompatibility is partial, a fraction of self-fertilization in a background of random mating yields a tangent vector 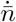 that cannot be distinguished from one produced by a shift in additive genic mortality together with viability selection against the heterozygote. (The symmetry analysis of App.A shows why a combination of the two is required.) The required shift depends on population state, however, so an incorrect generating model fitted initially to n will make the observed trajectory improbable over time.

Fig.7 illustrates the effect with a stark contrast using symmetry. Selectively neutral trajectories produced by admixtures of selfing and random mating (vertical lines in the figure), with total fecundity *γ*^R^+*γ*^S^ = 1, are initially fitted to models with fully random mating at *γ*^R^ =1 but both gametic and dominance viability selection. Trajectories with selection all sweep to fixation of either *g* = *aa* or *g* = *AA*, while neutral populations attract to a point between the Hardy-Weinberg and homozygous manifolds.

**FIG. 7:**
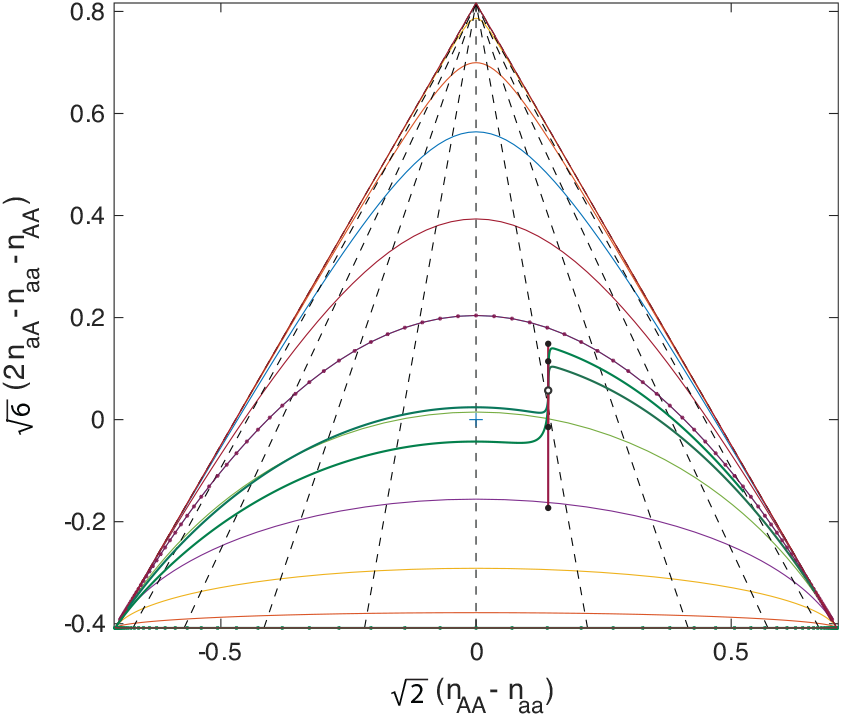
Each pair of purple and green trajectories compares two populations with *γ*^R^+*γ*^S^ = 1, one with neutral viability and nonzero *γ*^S^ (purple), the other with *γ*^S^ = 0 and both *μ*_||_ and *μ*_⊥_ non-zero (green). Values shown are *γ*^S^ ∈ {0.1, 0.25, 0.5, 0.75}. Filled black dots are approximating late-time fixed points of the neutral model.

Define the following symmetry-based coordinates for mortality, following the notations of Eq. (43):

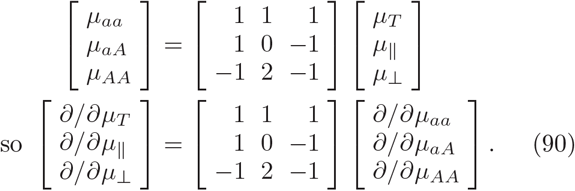

Fig.8 uses more realistic parameters, with additive selection at *μ*_||_ = –0.05, favoring allele *a* which is a minority in the initial condition, and *γ*^S^ = 0.05 from a low frequency of selfing. The only stark feature is an initial population significantly depleted in heterozygotes relative to the Hardy-Weinberg equilibrium, which we put in by hand to illustrate effects. A mis-specified model is fit using *γ*^S^ = 0 with over-dominance *μ*_⊥_ ≈ 0.021 and slightly shifted additive viability selection *μ*_║_ ≈ –0.058. Populations under the two models trace out nearly identical trajectories, but at different rates.

**FIG. 8:**
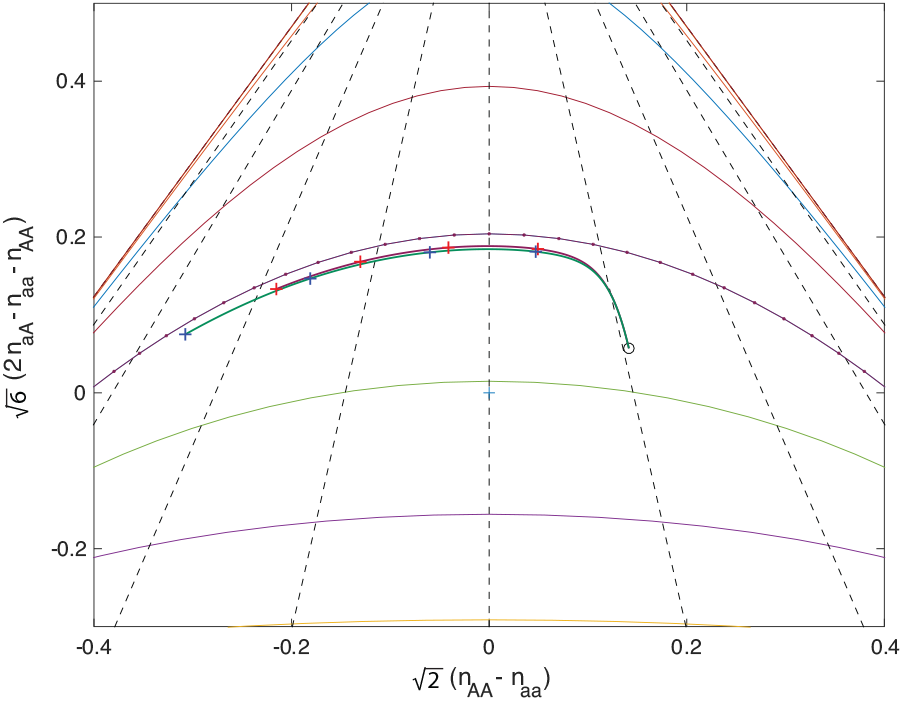
Two population models with the same initial tangent *ṅ*, one with *γ*S = 0.05 and *μ*_||_ = −0.05, the other with *γ*S = 0, *μ*⊥ ≈ 0.021, and *μ*_||_ ≈ −0.058. Markers (+) indicate generations 5, 10, 15, and 20 along each trajectory. The trajectory with overdominance runs toward fixation more quickly than the one that suppresses the heterozygote by selfing.

Fig.9 shows the integral information measure (80) and the integrand with the *γ*^S^ = 0.05 lifecycle as the null model. Information accumulates over the 20 generations shown mainly due to divergence in the sweep toward fixation of *a*.

**FIG. 9:**
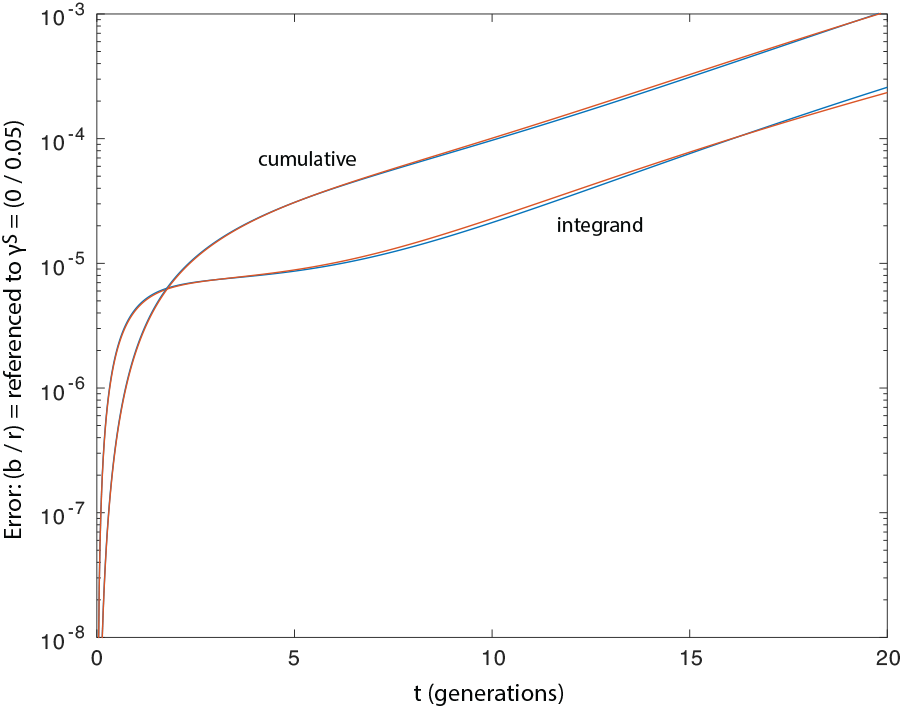
Information (80) in a trajectory generated by a model with γ^S^ = 0, relative to a null model at *γ*^S^ = 0.05 with which the trajectory agrees at *t* = 0, from Fig. 8. Integral and integrand are indicated.

### B. Contribution of lifecycle processes to total information

Fig.10 shows the information (86) (cumulative and information rate) of both models against the neutral model at the initial state 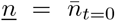. The rate of information input, initially high, falls rapidly over the first few generations while the population approaches the Hardy-Weinberg equilibrium, and only later rises gradually as it migrates within the attracting manifold.

**FIG. 10:**
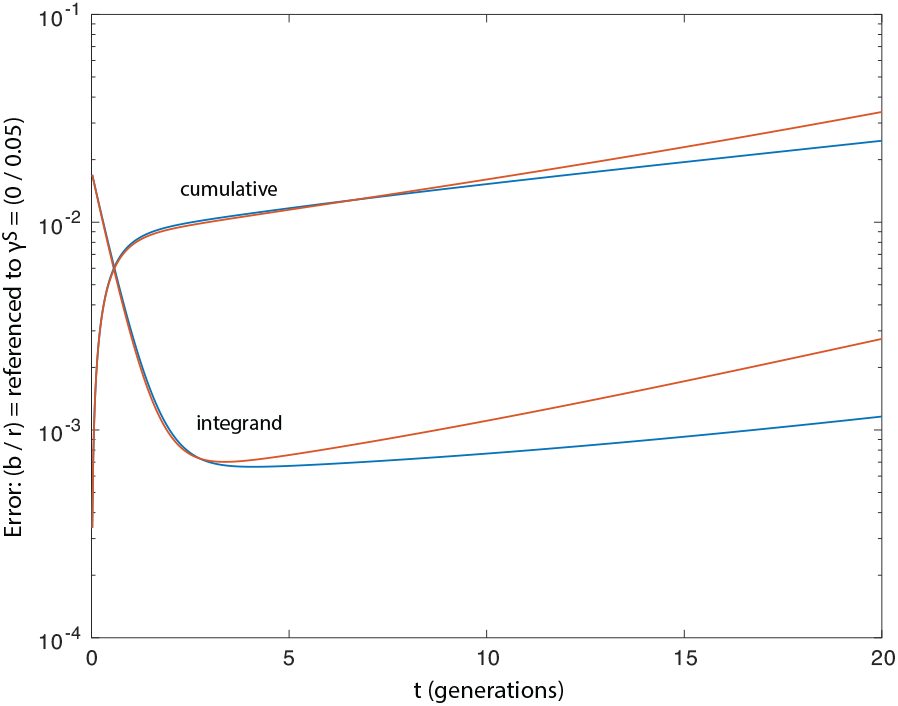
Fisher information measure obtained from Eq. (80) with the martingale as the null expectation for the trajectory. Again, whole trajectory error and “rate of information input” (integrand) shown versus the martingale reference model.

We study components of information inflow with the logarithmic derivatives 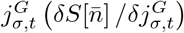 and 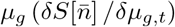; the sum of these over all currents equals 2×the total trajectory information. A combination of fecundity and mortality derivatives that governs the relative frequencies *n_g_*/*N* is

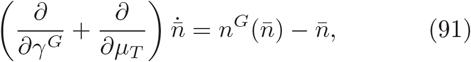

where the population structure 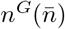 deposited through reproduction was introduced in Eq. (35).

In Fig.11, the current components for random mating are computed as 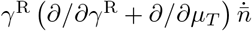 for the model with *γ*^S^ = 0.05, and 1 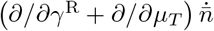 for *γ*^S^ = 0. The difference component from exchange of selfing for random mating at γ^S^ = 0.05, computed as 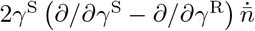, is compared to current from over-dominance 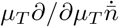 at *γ*^S^. Finally, the additive genic mortality currents are 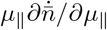 for the values in the two models.

**FIG. 11:**
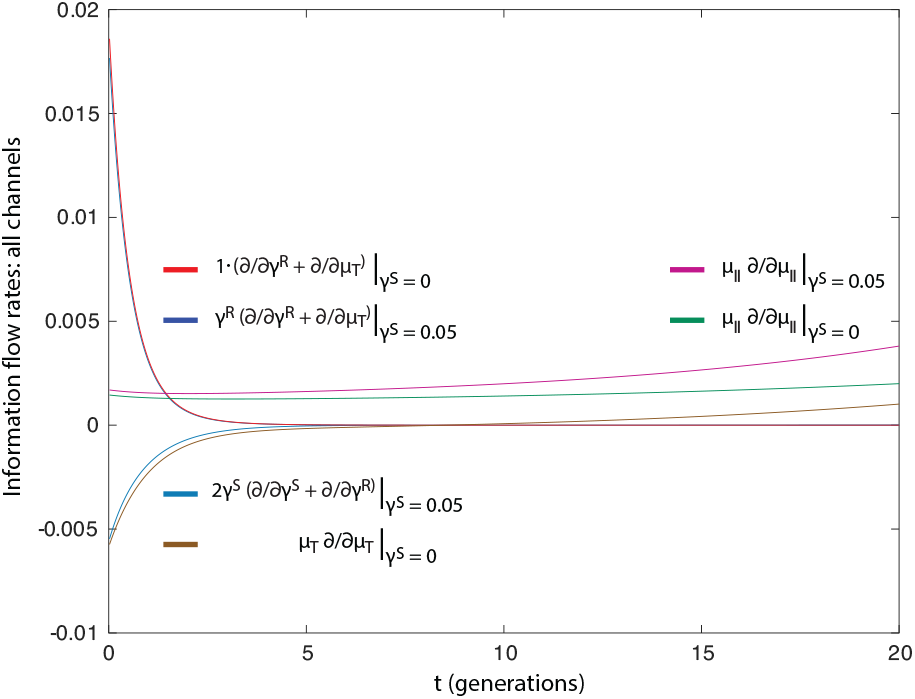
Information flows through three channels, for the two initially-confounded models of Fig. 8. Currents 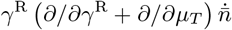 and 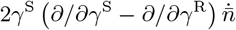 (labeled) in the model with *γ*^S^ = 0.05 and *μ*⊥ = 0 are compared to currents 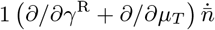 and 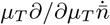 in the model with *γ*^S^ = 0 and *μ*_⊥_ ≈ 0.02. In both models, currents 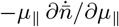 are compared. Random mating confers positive information flow because the current is in the direction toward the Hardy-Weinberg equilibrium, while selfing or excess heterozygote mortality are randomizing as their direction is toward the initial value *<u>n</u>*.

The figure shows that the *redistributing step* of fertilization – not expressible as selection in Price’s original scheme [1] – is responsible for the initially high and falling information input in Fig.10. The smaller currents from a partial substitution of selfing in one model, or increased mortality of heterozygotes in the other, enter as randomizing effects, pushing the population away from the Hardy-Weinberg equilibrium manifold and back toward its initial condition *<u>n</u>*. Information from additive mortality is smaller at both values of *μ*_||_ and increases with cumulative divergence of the population from the initial state.

## X. DISCUSSION

### A. Other applications

This paper has been concerned with showing why stoichiometric lifecycle representations are statistically needed, why they are a logical extension of principles already underlying population genetics, and what are their consequences for evolutionary concepts of causation and information. The methods themselves are easy to understand and use, and once recognized, they can immediately be applied to a range of problems. A few of those are noted below.

#### Epistasis and complex genotypes

Dominance and epistasis are routinely invoked together [10, 14, 17, 23] as the interaction effects that additive models handle poorly. The examples above treat only dominance. Epistasis can be handled in the same manner, and brings with it a reason to consider more complex combinatorial spaces of genotypes. Numerous standing problems such as the status of the chromosome as a unit of selection [34, 40, 45] could be well-systematized with such an approach.

#### Development

Population genetics conventionally compares alleles that alternate within fixed development and mating systems, with the latter falling outside the statistical study of evolution as change in frequencies. Sec.VI shows for the mating system how structural features of the lifecycle ordinarily left out of a frequency analysis are naturally absorbed in the covariance term of the Price equation on autocatalytic flows. The existence of development *at all*[6, 85] – the sorting of genetic interactions into a relatively conserved program carried in distributed form across many components making up the phenotype, and a variety of interchangeable modules subordinate to that program – is perhaps the most striking departure of biology to be explained, from an imagined primordial era of small loosely-organized RNA fragments [86, 87]. It would be desirable to capture the selective forces behind the emergence of these “algorithmic” dimensions of biology within the paradigm of evolutionary genetics itself.

#### Complex lifecycles

A two-phase lifecycle with ephemeral gametes is the simplest non-trivial generalization beyond replicators. Sexual reproduction together with multicellular development has opened up a host of much more complex and interesting lifecycles, particularly in the so-called “minor taxa”[63]. Even contrasts of monoecious and dioecious lifecycles in plants show the complex interplay of free-living stages and redistributive steps through which selection can act. It will be interesting to place the stoichiometric lifecycle models for the different known cases on a tree of their evolutionary relations, to learn whether lifecycle graphs decompose into natural modules corresponding to the developmental mechanisms enabling different stages.

#### Frequency-dependent effects

This paper has em-phasized the features of selection that cannot be subsumed within the abstraction of fitness, for which construction is a necessary addition. Numerous features that can be subsumed within fitness, but which involve dependence on other population members than the focal individual, are also well-handled with stoichiometric models. Most obvious among these are frequency-dependent selection effects, studied in stochastic stoichiometric models in [40]. The net effect of frequency dependence begs to be disaggregated into the fine structure of interaction within lifecycles, opening applications for a more diverse array of concepts from game theory [88] than those originally employed by Maynard Smith and Price [89, 90], which are particularly well-suited to generalizations to social evolution [91].

### B. A method for organizing and prioritizing summary statistics

While formally there is no difficulty in considering type spaces for whole genotypes, in practice low-order additive models are often the most that can be identified because there are too many genotypes for the statistical power available in regressions to make functional distinctions among them all. Heretofore, the solution has been to work out algorithmic traits of organisms such as their developmental programs or mating systems by other methods, leaving genetics to assign fitness differences to alleles within otherwise fixed systems.

The limitation of statistical power faced by the geneticist is also faced by selection itself [92]. Reinforcement learning, to which selection is equivalent [93], is capable of solving problems of only polynomial complexity, so the exponentially many variants of large genotypes cannot all have been assigned functional information through selection. There must be a principle of parsimony, and a domain of robust functional organization, within which only a subset of genetic variations carry selected information. An ideal evolutionary genetics would reflect the basis used by selection in nature, being neither too restrictive nor too inclusive.

The problem with writing a commitment to additive models into genetic methods is that, while parsimony may be a principle of selection, additivity generally is not. Additivity originates from considerations of symmetry, which parsimony in evolution need not observe.

Additive models relate to whole-genotype spaces as the first term in a direct-sum rather than a direct-product representation of genotypes. Let *i* ∈ 1,…, *N* index loci, and {*A_i_*}be a set of random variables for the alleles at each locus *i*. Then the random variable for a genotype may be written in direct-product form as a conjunction over loci, *g* = *A_i_* ∧ *A_2_* ∧ … ∧ *A_N_*. In the corresponding direct-sum decomposition, the basis elements are the disjunctions *g*^(1)^ = ∨_*i*_*A_i_* (the additive regressions), 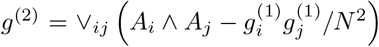 (excess second-order interactions), etc., and 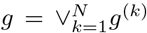. A way to reject most of the combinatorial diversity of genotypes while treating loci symmetrically is to restrict to low-order interactions in the direct sum.

Clearly a complex algorithm such as development or mating does not project into either a few factors in the direct product or a few summands in the direct sum, but rather cuts across either of these. Hypergraph generators for lifecycles may be understood as a formal system for representing evolutionary parsimony, which allows selective inclusion of interaction effects at multiple orders, within a uniform method of regression analysis.

### C. Managing combinatorial state spaces algebraically with rules

Regression is one type of infinitary-to-finitary map from properties of populations that are indefinitely (or even infinitely) larger than the rank of generating parameters on the hypergraph that they are used to estimate [55]. The generator itself, however, could also become indefinitely large – a case that would arise naturally and be desirable to consider if each species is a genotype – and such generators present their own problems of reduction and comprehensibility except in simple cases with very high symmetry.

Large structured type spaces in generators can remain tractable and comprehensible, however, if the species and reactions in them are produced by *rules*. An example of a rule for genotypes might be homologous recombination during meiosis: a pair of input chromosomes can be converted to any of a set of pairs of distinct output chromosomes, but reactions exist only to pairs that conserve sub-strings on either side of allowed crossover points. A single crossover rule automatically generates all recombination reactions in the hypergraph, so their construction and indexing can be done automatically by computer.

The derivation of indefinitely large types spaces in a generator by rules defines a second infinitary-to-finitary map, in which the finite range is the *rule algebra*[59, 94]. Rules specify equivalence classes of reactions, and from their algebra it is possible to extract the statistics of common features across multiple reactions, or to compute the interdependencies of reactions without solving for complete states of the system [95].

Rule-based modeling endows the species in the hyper-graph generator with a semantics: it is the properties of species that dictate which rules generate a reaction connecting them. A powerful example comes from the chemical reaction systems invoked at the beginning of this article [56, 57, 96]. What makes a molecule different from the list of its constituent atoms is the graph of their bonds (potentially with other properties as well). Graph rewriting rules, which act on fragmentary bond patterns rather than on entire molecules, formalize the notation of *reaction mechanisms*, and may be used to automatically generate indefinitely large reaction networks that are highly structured but much less symmetric than string representations of genotypes.

The Kappa programming language [59] was created to provide a platform to express rules of systems biology in an algebra of *site-graph* rewriting. Although originally conceived for intracellular processes such as protein binding, the same abstraction applies naturally to jointly-generated-phenotype concepts such as infectivity or virulence in host-pathogen interactions [23], gamete compatibility in fertilization, or interactions in ecosystems that make demes carriers of distributed information.

## XI. CONCLUSIONS

Seeking a general theory of selection in the organization of lifecycles rather than in the lineages of genes motivates the use of stoichiometric models as an alternative representation to Price’s corresponding sets. Lifecycle modeling generally precludes the categorical distinction between objects and relations that the replicator abstraction for genes takes as its starting assumption.

Relations in this context include any patterns that make an object assembled from components more than, or different in kind from, the list of its parts. Reliably establishing relations is the precondition to using information distributed across components to construct those phases of the lifecycle that are not made by direct copying. The goal in lifecycle modeling is to represent the conditions that make relations heritable, and the forms of selection that act through them as a result. Doing so is tantamount to keeping a part of the description of gene function explicit within the accounting procedures of genetics, which have conventionally been treated as external to it.

The concept of fitness has been strained for too long by an ambiguity of usage, between Darwin’s criterion of differential reproductive success (a lifecycle property) and Price’s definition (generalizing Fisher) from apportionment in corresponding sets, which generally cannot be made to span entire lifecycles. Replicators have taken on outsized importance in population genetics because they are the only abstraction restrictive enough to keep the two concepts of fitness from coming into conflict. The lack of a categorical resolution to problems of multilevel selection helps to identify the areas of conflict between fitness concepts, but the source of these problems is the inadequacy of the replicator abstraction and the insufficiency of fitness itself.

Stoichiometric modeling of lifecycles provides a systematic program to construct a suite of additional summary statistics beyond fitness describing the constructive steps in reproduction that fall outside the scope of replication. The Price equation in this broader suite of summary statistics moves several sources of change, including redistributive effects, out of the so-called “environment” term and into the covariance, which then takes on much more of the causal association with selection that Fisher’s Fundamental Theorem and its relatives were meant to possess.

Regressions on flows do this in a different and more complete way than abstractions such as jointly-generated phenotypes, because flows are inherently *concurrent processes*, whereas phenotypes are properties attached to states. Complex objects, which for some purposes may be viewed as collections, exist in the stoichiometric representation along with the sets that mediate flows, but the carriage of evolutionary memory and the control of evolutionary dynamics are more clearly distinguished and more flexible in the hypergraph than in the ordinary graphs for Price’s corresponding sets.

Extensive probabilistic and statistical methods exist to study stoichiometric population processes. Among these are large-deviation methods from which we can define a natural measures of the information in a population trajectory not yet incorporated in a generating model. Decomposition of this information into the contributions from flows defined on lifecycle graphs delivers the relation of selection to entropy that Price sought. Construction within lifecycles is amenable to a variety of automated and rule-based methods, many of them already well-developed in systems biology and other fields. With these we may import much more of the known complexity of biology systematically into the concepts and methods of genetics.

## Appendix A: Mating invariants, exponential families, and dualities

This appendix contains three derivations for the biallelic sexual reproduction system to provide a foundation for the analysis of examples in the main text. All are elementary, and some employ constructions dating back to Fisher. They are derived together here to fix notation and to recall features of regression models that appear in solutions in the text.

First, the manifolds of fixed points under the two mating systems are characterized, as the organizing centers for the dynamics. Second, the duality is explained between additive selection, and the Hardy-Weinberg family where additivity reduces to proper allelic fitness. Third the exponential families generated by the components of viability selection are related to the families generated by the fertilization systems in the absence of selection.

### 1. Invariant families for the two fertilization systems

Both the discrete-generation maps (1, 2) and the continuous-time counterparts (3, 4) are characterized by the fixed points of their reproduction models, which are also the asymptotic population states in the absence of fitness differences. Taking *n_a_* and *n_A_* as defined in Eq. (15) at 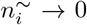, and 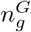 as introduced in Eq. (35), these fixed point solutions are respectively

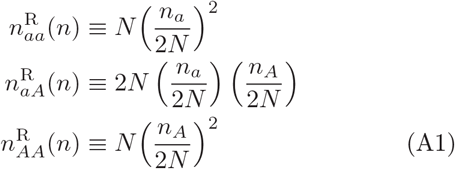

for random mating, and

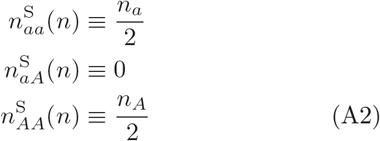

for selfing.

As must be the case for a purely redistributive process, allele counts are preserved under either fertilization map:

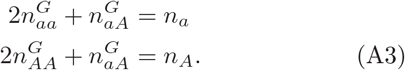

Therefore the preimages of either of the maps (A1, A2) in the projection *n/N* are vertical lines, as shown in Fig. 12. Likewise, the contributions *n^G^*(*n*) – *n* to *d*(*n/N*) /*dt* in either of (3, 4) at neutral gametogenesis fall within these vertical loci.

**FIG. 12:**
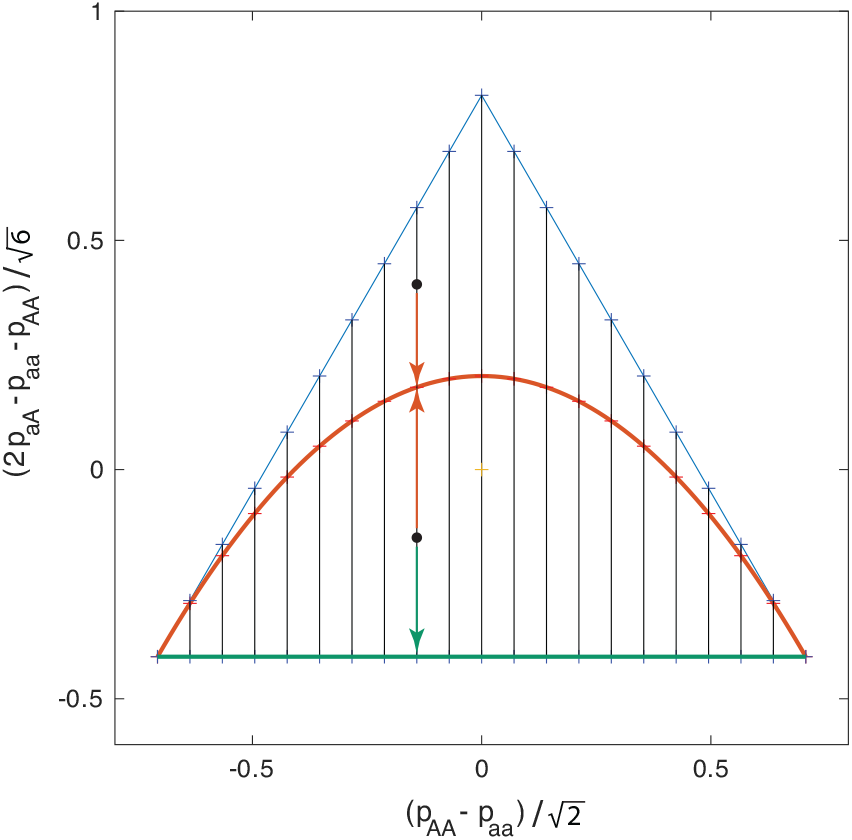
The simplex of vectors (*n_g_/N*) viewed from its normal in the positive orthant of R3. Hardy-Weinberg equilibria (A1) (heavy red) with 20 uniformly-spaced values of *p_a_* ∈ [0, 1] marked with symbols. Invariant manifold for selfing (A2) (bold green), is the lower boundary. Pre-images of fixed nG are vertical black lines. Components *n_G_* (*n*) – *n* of neutral trajectories under either mating system are linear vertical attractors from states *n* (two black dots as examples) toward the Hardy-Weinberg or homozygous (*n_aA_* = 0) boundary, (colored arrows).

Both character averages 〈*χ*〉 and the fixed points (A1, A2) take simple forms in terms of the normalized measure *n/N*, so denote

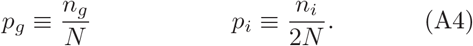

Fisher’s spherical coordinates 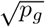 for distribution *p*(see [97]) can be written

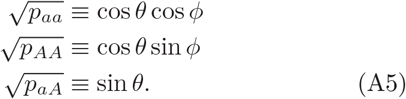

The invariant manifold (A2) for selfing corresponds to *θ* = 0.

The Hardy-Weinberg equilibria (A1) satisfy 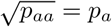 and 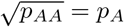, or in the polar coordinates (A5)

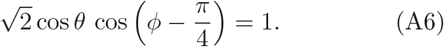

These points form a circular arc which is the intersection of a vertical plane with the Fisher sphere, sharing the endpoints *p_aa_* = 1 and *p_AA_* = 1 with the *θ* = 0 invariant family for selfing. Let *ν* ∈ [0, *π*] be an angle from the horizon in the vertical plane, referenced to the midpoint of the chord, *p_aa_* = *p_AA_* = 1/2. Then

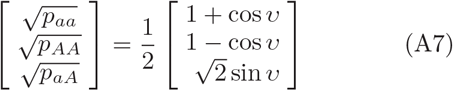

expresses the family in polar coordinates on a circle of radius 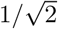 within the plane.

### 2. Duality of additive fitness and Hardy-Weinberg equilibrium

When additive models are used to project fitness characteristics of diploids onto a basis of haplotypes, they may be mis-specified because either the diploid population state or the fitness function has information that the coefficients in the additive decomposition cannot represent. An additive model may fail to characterize the dynamics of the alleles that it is nominally defined to describe, or the dynamics of heterozygosity that by construction it omits. Within the space of population states (*p_g_*), however, the Hardy-Weinberg manifold is an exponential family generated by additive fitness in the absence of selection on heterozygosity, where the regression coefficients become proper allelic fitnesses and the dynamics reduces to that of the one-dimensional distribution (*p_i_*).

#### a. Additive models of pure viability selection on genotypes

To separate the diploid fitness problem from the mating system, take the coefficient *γ* that scales 〈Δ*^G^χ*〉 in the Price equation (38) to zero relative to (*μ_g_*). The rate equations for the population state then become

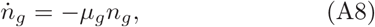

with *μ_g_* constants. State-dependent effective fitness coefficients *μ_i_*(*n*) on alleles are defined from

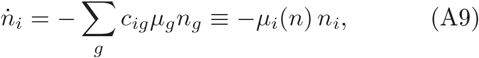

in terms of the stoichiometric coefficients *c_ig_* from Eq. (14). Total mortality is denoted as 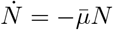, with

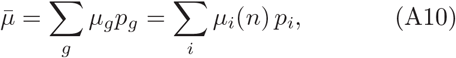

giving the rate equations for genotype or allele fractions

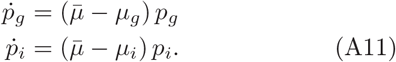

The changes in allele fractions relate to the *μ_i_* (whether constant or state-dependent) as

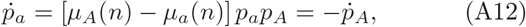

or equivalently, for the tangent vector to (log *p_i_*),

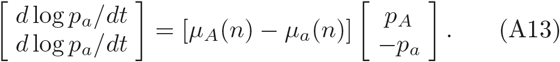

An additive regression model of the trajectory *p* generated by (*μ_g_*) will generate a trajectory *p*^add^, with *p*^add^ = *p* as initial condition and 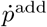 as close as possible to 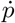. Fitness coefficients in the so-called “*α*-representation” (see [14]) are given by

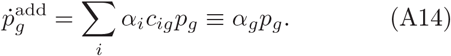

Although the regression coefficients *α_i_* will inevitably be state dependent in the general case, they are the independent parameters in terms of which diploid coefficients *α_g_* are defined. Because only *p* rather than *n* is modeled, we must set 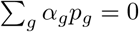, implying that for some λ

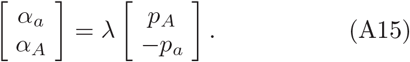

The conventional least-squared error criterion to fit *α* has the same form as the error rate in the large-deviation probability (80) for paths:

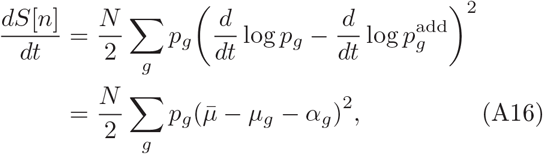

only evaluated in the background of a neutral model 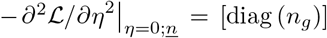 rather than in the background of either of the generative models for *p* or *p*^add^.

Evaluating Eq. (A16) along the ray (A15) and then minimizing over λ gives

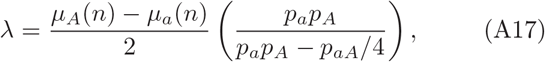

so the best-fit additive coefficients relate to the actual tangent (A13) to (log *p_i_*) as

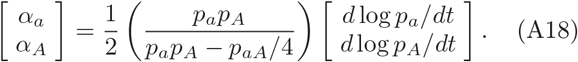

The first observation is that within the Hardy-Weinberg family (A1), where *p_aA_* = 2*p_a_p_A_*, the coefficient of proportionality equals unity; everywhere else it differs by a state-dependent multiplier. Thus, independent of whether the generating model *μ* responsible for d log *p_i_*/*dt* is additive, the *α* coefficients are only descriptively sufficient (see Lewontin [34] for the origin of dynamical and explanatory sufficiency) when fit to Hardy-Weinberg starting populations.

Descriptive sufficiency of the additive model is independent of whether the model also omits by construction the intensity of selection on heterozygosity in the generating process. To characterize the error from the latter, we decompose the definitions (A9) into the relation

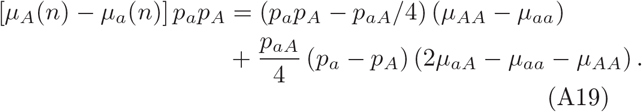

Eq. (A15) and Eq. (A17) together then give

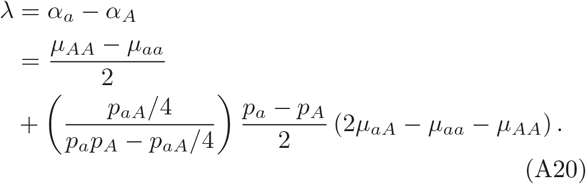

If 2*μ_aA_* – *μ_aa_* – *μ_AA_* = 0, the difference of the additive coefficients (*α_a_ – α_A_*) is constant (their sum is fixed by an independent constraint), and recovers entirely the mortality differential in the generative process. In that sense the additive model is dynamically sufficient even while failing to coincide with an actual constant gametic fitness in the general case by Eq. (A18).

The regression coefficients *α_i_* more generally are state dependent. In the genotype basis *g*, the relation (A20) implies that

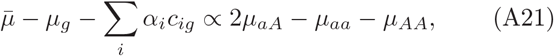

with a state-dependent coefficient of proportionality that we do not write down. It then follows that the minimum residual error (A16) is nonzero with

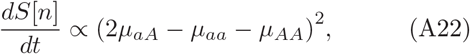

again with state-dependent coefficient of proportionality.

The residual error function can be decomposed into a sum of terms characterizing descriptive sufficiency within the basis *i* and a difference of covariances between the *g* and *i* bases. Because the *α*-representation is additive, the definition (A14) can be re-arranged to

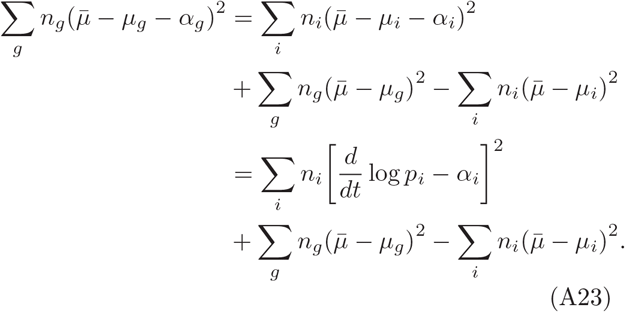

In the Hardy-Weinberg manifold where Eq. (A18) gives *d*log*p_I_*/*dt* = *α_I_*, the residual error (A22) comes from the excess variance of mortality in basis *g* due to selection on heterozygosity, over the variance in the basis i for alleles.

#### b. The Hardy-Weinberg family is the dual exponential family to additive fitness

Now restore *γ* > 0 to consider viability selection in the presence of reproduction. For either fertilization system, within its own invariant manifold *n^G^*(*n*) – *n* = 0, and there the rate equations (A8) become

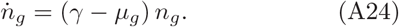

The constant *γ* changes only 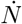 from the previous case, and all subsequent analysis proceeds as above.

The homozygous family *p_aA_* = 0 is preserved by selfing, but there *α* accounts for only 1/2 the tangent vector *d*log*p_i_*/*dt* in Eq. (A18). In contrast, the Hardy-Weinberg family, preserved by random mating, is identified with additive fitness by the condition of descriptive sufficiency on initial p, while additive fitness is identified by the condition (A23) of zero excess variance as the generating process that leaves *p* within that manifold, resulting in constant *α_a_* – *α_A_* = *μ_A_* – *μ_a_* that is a proper allelic fitness difference.

### 3. The exponential family in additive and dominance coordinates

The generating coefficients 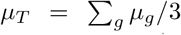, *μ*_║_ ≡ (*μ_aa_ – μ_AA_*)/2, and *μ*_⊥_ ≡ (2*μ_aA_ – μ_aa_ – μ_AA_*)/6 introduced in Eq. (90) are dual coordinates to an exponential family in variables *N* and *p*. Fig. 13 shows the projections of these families in the basis set *∂*/*∂*_*μ*║_ and *∂*/*∂*_*μ*⊥_.

**FIG. 13:**
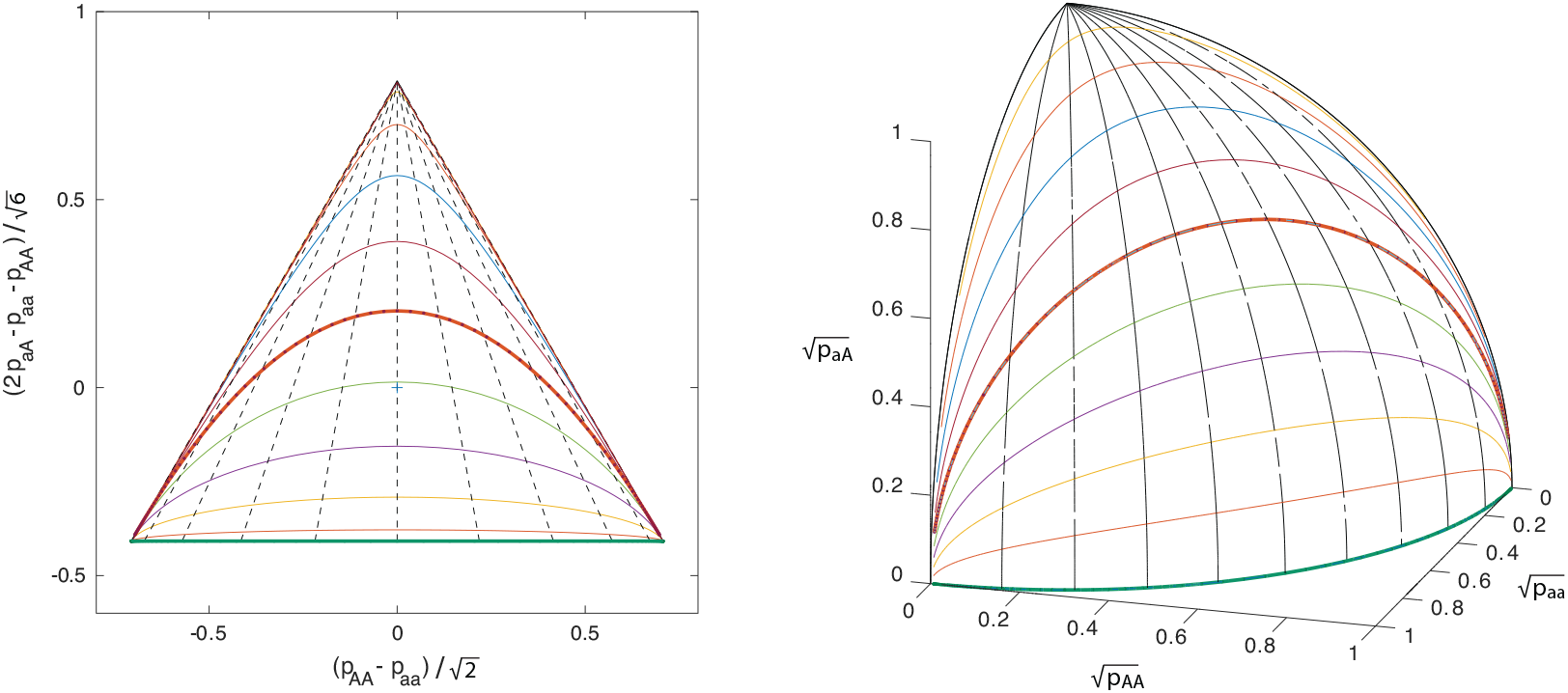
Additive exponential families and dominance-driven trajectories as coordinates in the simplex and in Fisher’s spherical coordinates (A5). The H-W family and the selfing-invariant family *p_aA_* = 0 are marked. Allele neutrality but dominance 2*μ_aA_* – *μ_aa_* – *μ_AA_* ≠ 0 preserves *p_aa_/p_AA_*, leading to azimuthal planar sections on the Fisher sphere.

Latitudes are generated by additive diploid fitness. Longitudes, generated by selection on heterozygosity (dominance), are sections of fixed *p_aa_*/*p_AA_*. The transformation of the vertical basis vectors (a mixture family [97]) generated by *∂*/*∂γ* in Fig. 12 into coordinates (*∂*/*∂*_*μ*║_, *∂*/*∂*_*μ*⊥_ is used to define the confounded models at a single value of 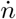 in Sec.IX.

## Appendix B: Current component decomposition for models of asymmetric gametogenesis

This appendix provides supplemental algebraic forms for the regression coefficients corresponding to the break-down in Equations (21,24), for the two cases of uniform asymmetric gametogenesis in Sec.VB and meiotic drive in Sec. (VD).

### 1. The uniform asymmetric gametogenesis case

A set of current coefficients for the case of uniform gametogenesis, respecting the scaling (20) with *N*, for random mating, is given by

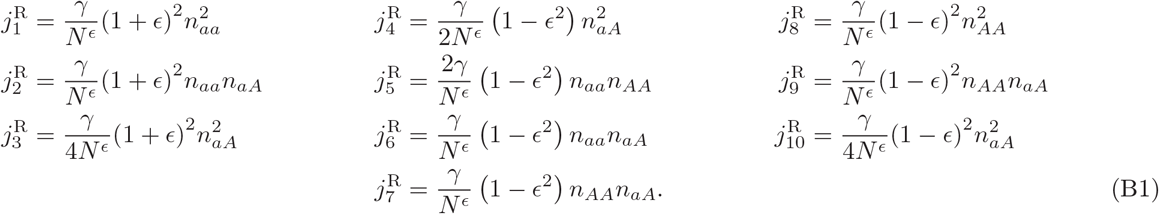

The corresponding currents for self-fertilization are

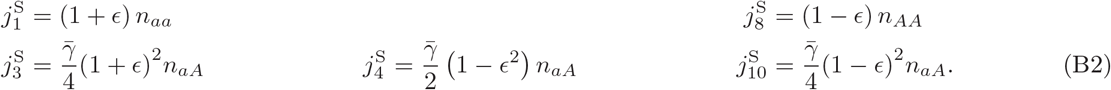

### 2. The meiotic drive case

A set of current coefficients for the case of meiotic drive, respecting the scaling (20) with *N*, for random mating, is given by

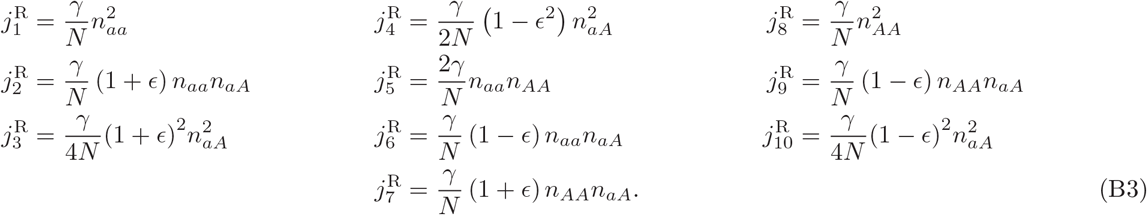

The corresponding currents for self-fertilization are

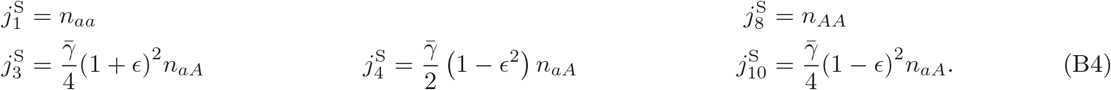

## Appendix C: Two-field functional integral methods for stochastic population processes

A two-step process is used to convert generating functions, as analytic functions of complex variables, into the extended-time representation of 2-field functional integrals. First the formal power series properties of the MGF are separated from its behavior as an analytic function by replacing complex arguments and their partial derivatives with a formal algebra of raising and lowering operators. Second an overcomplete basis of eigenstates of the raising and lowering operators is used to furnish a representation of unity in the Hilbert space of MGFs, permitting the conversation of the formal quadrature of the Liouville equation to a functional integral on the eigen-values as field variables of integration. More thorough discussion surrounding these steps for population processes of varying complexity may be found in [39, 40].

### 1. The Doi algebra for moment-generating functions as formal power series

The operator algebra for MGFs is due to Masao Doi [73, 74]. One studies the MGF as a formal power series by replacing the complex arguments of Eq.(63) with vectors of raising and lowering operators 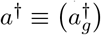, *a* ≡ (*a_g_*) under the map

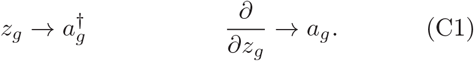

The feature of analytic functions inherited by the formal operators is the commutation relation 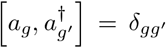, where *δ_gg′_* is the Kronecker *δ* symbol.

The space of MGFs is constructed as a Hilbert space with inner product by introducing dual “ground” states |0) and (0| which correspond, in the space of analytic functions, to the mapping

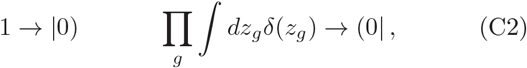

where *δ*(*z_g_*) is the Dirac *δ* distribution and the integral in Eq. (C2) is to be applied to any analytic function. Under the mappings (C1,C2) the MGF (63) becomes a formal state

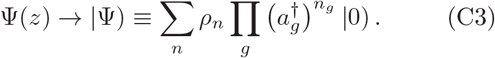

All states are normalized under the inner product with the *Glauber state* 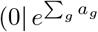, which extracts the trace of the probability distribution

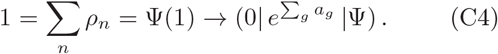

The representation in analytic functions can be recovered through an inner product with a generalization of the Glauber norm,

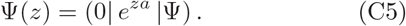

The Liouville equation (64) then takes the form

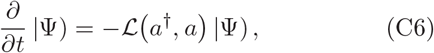

in terms of the function 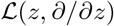 under the argument substitution (C1). Eq. (C6) may be formally reduced to the quadrature for |Ψ) at any time *T* >0 as

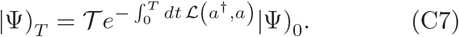

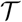 denotes the time-ordering operator on exponentials, with 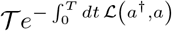 understood as a time-ordered matrix product, over small intervals *dt*, of 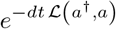.

### 2. The Peliti coherent-state basis and functional integral

A generating functional and time-dependent correlations functions can be constructed by placing insertions in the time-ordered quadrature (C7). Peliti [75, 76] intro-duced the step of inserting a representation of unity be) tween small increments of time evolution under 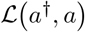 to convert the operator quadrature to a functional integral.

For a vector *ξ* ≡ (*ξ_g_*) of complex coefficients and *ξ*^†^ its Hermitian conjugate (complex conjugate and transpose), the state |*ξ*) and its dual projection operator (*ξ*^†^| are eigenstates of the lowering and raising operators respectively:

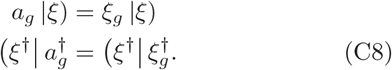

|*ξ*) are termed *coherent states*, and are the generating functions for products of Poisson distributions over each *n_g_* with parameter *ξ_g_*. With suitable normalization, it can be shown [39, 40] that dual coherent states have inner product

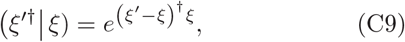

and the following integral is a representation of the identity operator in the Hilbert space of generating functions:

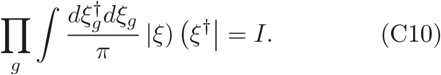

Inserting the Peliti representation of unity (C10) at intervals *dt* in Eq. (C7) gives the functional integral

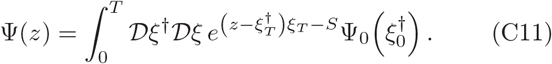

The functional measure is defined as the limit of a skele-tonized product measure

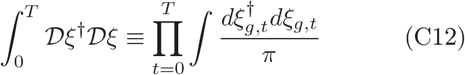

as *dt* → 0. As long as 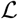 is written in what is termed *normal order*, with all factors of *a_g_* to the left of any factor of 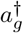, the action *S* in Eq. (C11) can be computed from the eigenvalue property (C8) and the inner product (C9) to be

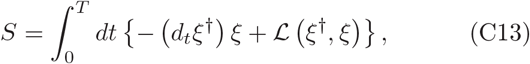

with 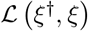 obtained from the form in Eq. (C6) by the argument substitution *a*^†^ → *ξ^†^, a* → *ξ* from Eq. (C8).

The natural observable *n_g_* for which to study correlations is produced by insertion of the product of Doi operators 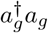 in expectation values, and is therefore given in the field integral (C11) by insertions of the product 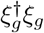. It is possible to work with the number variable as an elementary field by writing the coherent-state field variables in the functional measure (C12)

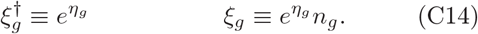

The change of variables (C14), which is a canonical transformation [98] in the Hamiltonian system generated by 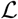, converts the functional integral (C11) and action (C13) into the forms (65) and (66) in the main text.

## Footnotes

1 These framings, which concern conventional biological individuals or sets or parts of individuals, are to be contrasted with work on channels for evolutionary information flow through systems that are not individuals subject to vertical descent, as in the work on Niche Construction [8].

2 Here, we follow Ewens and Lessard [14] in restricting FTNS to apply to additive genetic variance in diploid models, and use the term “growth rate equation” for more general covariance expressions.

3 As noted, Niche Construction [8] seeks to capture memory mechanisms affecting evolution, conceived of as operating outside of and in parallel to Mendelian genetic heredity, though coupled to it through genes’ determination of traits. Flack *et al*. [18] have argued for a much more inclusive paradigm that they term *construction dynamics*, encompassing the many ways in which distributed information determines the biological or social architectures behind abstractions such as the genotype-to-phenotype map. While invoking Niche Construction in early presentations [19], and mostly emphasizing time-scale or information hierarchies as examples, construction dynamics refers equally to the mechanisms determined by distributed information that are the examples in this paper.

4 The distinction is between a naïve concept of reductionism, identifying an object only with the list of its components, and the concept of reductionism actually used in any mature science, in which properties of building blocks exert constraints on the kinds of assemblages that can be created from them, which if they are understood can simplify the problem of characterizing the assemblages.

5 This is so, notwithstanding the problems with such interpretations [14, 22, 24] noted above.

6 A multiset is a sample of individual objects in which multiple objects may be of the same type [57].

7 In more elaborate models, the null object can also be a source for introduction of individuals to the population, modeling phenomena such as migration as in [58].

8 Note that, while the figures represent these objects with doubly-bipartite simple graphs (two kinds of nodes and two kinds of edges), the distinct meaning of the different classes of nodes and edges with respect to the Markov state and dynamics signals that the underlying mathematical object is the hypergraph.

9 The one relation *γ/κN* → 0 is sufficient to produce comparable deterministic rate equations for the two fertilization systems. In a stochastic treatment, more care would need to be given to the particular scaling of *κ* relative to *N*, to reflect the organism-level physiology of selfing in relation to the deme-level statistics of gamete sampling for random mating.

10 “Competition” is used broadly to refer to changes that are in tension among objects, which may or may not be of the same types. Formalizations of the term encounter difficulties, however, for dissimilar objects. In game theory [65], competition is a relation among peers under rules of the game. If the population-genetic definition of fitness is taken to formalize a notion of competition in evolution, then that formalization is likewise limited to objects of equivalent type, for which fitness is commensurable. An extension of population-genetic ideas to regression coefficients beyond fitness and heritability provides a way to capture a larger scope of the pre-formal meaning without retreating to metaphor. A similar problem affects “levels”. The stoichiometric representation admits objects of arbitrary types, some sets of which may be interconvertible to others. It is only with the addition of either constraints or further interpretations that objects of different kinds are seen to relate to each other in “levels”.

11 Such a large value of ∈ is chosen to make the effect, which is 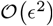, visible in the figure. Realistically such an invading haplotype would represent a qualitative change in the mating system. For more typical values of ∈ of a few percent, the consequent differences for change in population structure would be expressed over hundreds rather than tens of generations.

12 The sign used here to refer to refer to relative entropy coincides with that for Shannon entropy on the uniform measure. It is common, where the sign convention is clear, to find the Kullback-Leibler divergence identified as relative entropy.For introduction and definitions of terms see [79].

13 For introduction and definitions of terms see [79].

14 This term is adopted from the appellation for the same construction in *importance sampling* [80], to which large-deviation methods are closely related [78, 81].

15 A more formal treatment of what is meant by simplicity is given in [39]. Generalizations with concurrent firing of multiples of reactions can readily be defined in the operator algebra, at the cost of more complexity.

